# Nanobodies restore stability to cancer-associated mutants of tumor suppressor protein p16^INK4a^

**DOI:** 10.1101/2021.07.01.450670

**Authors:** Owen Burbidge, Martyna W. Pastok, Samantha L. Hodder, Grasilda Zenkevičiūtė, Martin E. M. Noble, Jane A. Endicott, Laura Itzhaki

## Abstract

We describe the generation and characterization of camelid single-domain antibodies (nanobodies) raised against tumor suppressor protein p16^INK4a^ (p16). p16 plays a critical role in the cell cycle by inhibiting cyclin-dependent kinases CDK4 and CDK6, and it is inactivated in sporadic and familial cancers. The majority of the p16 missense mutations cause loss of function by destabilizing the protein’s structure. We show that the nanobodies bind p16 with nanomolar affinities and restore the stability of a range of different cancer-associated p16 mutations located at sites throughout the protein. The nanobodies also bind and stabilize p16 in a cellular setting. The crystal structure of a nanobody-p16 complex reveals that the nanobody binds to the opposite face of p16 to the CDK-binding interface permitting formation of a ternary complex. These findings indicate that nanobodies could be used as pharmacological chaperones to determine the consequences of restoring the function of p16 in the cell.

**Highlights:**

- Describes a nanobody (single-domain antibody) capable of binding tumor suppressor protein p16
- Nanobody binding can stabilise wild-type and cancer-associated p16 mutants
- Nanobody-p16 crystal structure reveals why nanobody binding is compatible with p16-CDK6 interaction

## Introduction

The ankyrin-repeat protein p16^INK4a^ (p16) is a tumor suppressor that plays a critical role in regulation of the cell cycle by binding and inhibiting cyclin-dependent kinases 4 and 6 (CDK4 and CDK6) (Kamb et al., 1994; Morgan, 2007; Serrano et al., 1993), thereby blocking pRb phosphorylation and entry into S phase (Koh et al., 1995; Lukas et al., 1995; Medema et al., 1995). Gene deletion (Liu et al., 1995), transcriptional silencing (Merlo et al., 1995) or point mutations (Harinck et al., 2012; Kamb et al., 1994; Koh et al., 1995; Parry and Peters, 1996; Poi et al., 2001; Ranade et al., 1995) of the *CDKN2A* gene that encodes p16 result in loss of function of the protein, thereby driving tumorigenesis (Liggett and Sidransky, 1998; Sherr et al., 2016). p16 is a highly flexible (Byeon et al., 1998), protein of marginal thermal stability (Tang et al., 1999) and consequently many of the cancer-associated point mutations severely compromise the overall protein fold, resulting in loss of binding and/or an increased propensity to misfold or aggregate (Boice and Fairman, 1996; Byeon et al., 1998; Tevelev et al., 1996; Yang et al., 1995; Yuan et al., 1999; Zhang and Peng, 1996). Previous studies have shown that second-site mutations in similarly unstable proteins, such as the tumor suppressor protein p53, can stabilize the native structure and counteract these effects (Joerger et al., 2005; Nikolova, 2000; Otsuka et al., 2007). The incorporation of mutations at three sites in p16, designed to increase the stability of the protein, was found to suppress a range of cancer-related mutations (Cammett et al., 2003). Taken together, these findings suggest that stabilization of p16 by exogenous ligands - i.e. pharmacological chaperones - could be a therapeutic strategy to restore its stability and thereby rescue its activity. Indeed, given that the vast majority of disease-associated missense mutations are predicted to cause loss of function indirectly by destabilizing protein structures rather than directly by disrupting residues in catalytic or other functional sites (Landon et al., 2009), pharmacological chaperones to aid the folding of mutant variants could be applicable to many diseases (Beck, 2017; Maurer et al., 2018; Okiyoneda et al., 2013; Zawacka-Pankau and Selivanova, 2015). (Liu et al., 1995), transcriptional silencing (Merlo et al., 1995) or point mutations (Harinck et al., 2012; Kamb et al., 1994; Koh et al., 1995; Parry and Peters, 1996; Poi et al., 2001; Ranade et al., 1995) of the *CDKN2A* gene encoding p16 result in loss of function of the protein, thereby driving tumorigenesis (Liggett and Sidransky, 1998; Sherr et al., 2016). p16 is a highly flexible (Byeon et al., 1998), thermodynamically unstable protein (Tang et al., 1999) and consequently many of the cancer-associated point mutations severely compromise the overall protein fold, resulting in loss of binding and/or an increased propensity to misfold or aggregate (Boice and Fairman, 1996; Byeon et al., 1998; Tevelev et al., 1996; Yang et al., 1995; Yuan et al., 1999; Zhang and Peng, 1996). Previous studies have shown that second-site mutations in similarly unstable proteins such as the tumor suppressor protein p53 can stabilize the native structure and counteract these effects (Joerger et al., 2005; Nikolova, 2000; Otsuka et al., 2007). The incorporation of mutations at three sites in p16, designed to increase the stability of the protein, was found to suppress a range of cancer-related mutations (Cammett et al., 2003). Taken together, these findings suggest that stabilization of p16 using exogenous ligands - i.e. pharmacological chaperones - could be a therapeutic strategy to restore its stability and thereby rescue its activity. Indeed, given that the vast majority of disease-associated missense mutations are predicted to cause loss of function indirectly by destabilizing protein structures rather than directly by disrupting residues in catalytic or other functional sites (Landon et al., 2009), pharmacological chaperones to aid the folding of mutant variants should be applicable to many diseases (Beck, 2017; Maurer et al., 2018; Okiyoneda et al., 2013; Zawacka-Pankau and Selivanova, 2015).

Single-domain antibodies derived from naturally occurring camelid heavy-chain antibodies (nanobodies) have several advantages over conventional antibodies (Gettemans and Dobbelaer, 2021; Hamers-Casterman et al., 1993). Their ability to bind non-conventional epitopes with high affinities (Stijlemans et al., 2004), as well as their high thermodynamic stability (Dumoulin et al., 2002) and their small size, present many benefits for biotechnological and clinical applications (Hoey et al., 2019; Jovčevska and Muyldermans, 2020). Recently, nanobodies have been used to stabilize unstable proteins (Loris et al., 2003) and dynamic protein complexes as an aid to X-ray crystallography (Che et al., 2018; García-Nafría and Tate, 2019; Nordeen et al., 2020; Tereshko et al., 2008). Single-domain antibodies have also been shown to stabilize the native state of a disease-causing mutant of human lysozyme thereby inhibiting lysozyme-amyloid formation (Dumoulin et al., 2003; Pain et al., 2015). Similar anti-amyloid effects have been observed for nanobodies targeting alpha-synuclein (Genst et al., 2010) and huntingtin protein (Messer and Butler, 2020; Schiefner et al., 2011).

We describe the characterization of a set of anti-p16 nanobodies and demonstrate that two have a profound effect on p16 stability. Nanobodies capable of stabilising p16 also stabilized a range of different cancer-associated p16 mutants located at sites throughout the p16 structure, restoring stability to or beyond wild-type levels for some of the mutants tested. We present the crystal structure of p16 in complex with the more stabilizing of these two nanobodies, revealing that the nanobody binds on the opposite face of p16 to the face involved in binding to CDKs. We demonstrate formation of complexes containing nanobody, p16 and CDK6 in solution and in cells. This nanobody has the potential to be exploited as a pharmacological chaperone to stabilize and restore the cellular functions of cancer-associated mutant p16.

## Results and Discussion

### Generation and selection of the anti-p16 nanobody library: antigen production and characterization

The quality of an immune library to be used for selections by display techniques is related to the quality of the antigen administered to the animal and the size of the immune response generated against it. The incorporation of tags and modifications to previously reported p16 expression and purification protocols resulted in a significant increase in the yield, purity and homogeneity of the p16 proteins (StarMethods). Three point mutations (W15D, L37S and L121R), previously shown to stabilize p16, were used (both singly and in combination) to generate more stable p16 species (Cammett et al., 2003). p16 proteins (mutants and wild-type p16 N- and C-terminally Avi-tagged (p16-NTA and p16-CTA respectively)) were subsequently expressed and purified from recombinant *E. coli* cells. After *Llama glama* immunisation, two immune libraries (libraries 173 and 174 raised against wild type p16 and double mutant p16^L37S/L121R^ respectively) were cloned into a phage display vector pMESy4 and transformed into TG1 *E. coli*. After phage library rescue, titration of the precipitated phage for each library revealed a phage input for subsequent biopanning of 4.2 x 10^13^ for the wild-type library (173) and 6.2 x 10^13^ for the stabilized p16 library (174).

To identify a diverse range of different nanobodies that bind to p16 at the expense of binding affinity, the first round of nanobody phage display was performed using a high antigen concentration across six different biopanning strategies (StarMethods). Briefly, these strategies utilised solid phase coated p16 or p16^L37S/L121R^ on immunosorp plates either individually or bound to CDK6 or employed capture of p16 either by solid phase coated CDK6 or through neutravidin capture of the biotinylated AviTag. Each library was panned using these strategies before non-specific elution using trypsin to remove bound phage. Titration of the eluted phage following output library rescue revealed a large enrichment in nanobodies from Library 174 with minimal amounts of enrichment for the wild-type Library 173. A second modified round of phage display was then performed for Library 174 in which the antigen concentration was dropped to try and increase enrichment of high-affinity nanobodies. However, the antigen concentration for Library 173 was kept the same in this second round so as not to lose nanobody diversity. From each biopanning strategy, several unique clones were selected. Monoclonal phage were produced for each and then subsequently tested for binding to p16 by ELISA and sequenced. For most biopanning strategies using the double mutant library 174, positive phage binders were observed. Slightly fewer were seen for the wild-type library 173, with very few binders being found for the CDK6 capture of p16.

Positive binders were grouped by CDR3 sequence similarity into a number of families. Although family members shared similarities in sequence for CDR3, diversity was seen within other regions of the protein including point mutations within framework regions and differences in CDR1 and CDR2 loop lengths. Individual point mutations within the CDR regions may be attributed to somatic hypermutation and affinity maturation of the nanobodies. Six large families dominated the phage display output (**Figure 1A, Figure S1**) and made up over 60% of the nanobodies generated. Sixteen smaller families were also distinguished that consisted of only a few nanobody clones. Each of these families exhibited large sequence diversity in their CDR regions (**Figure 1A, B**). Twenty representative nanobodies were selected at random. Of these 19 were expressed at levels sufficient to enable further characterisation.

**Figure 1.**
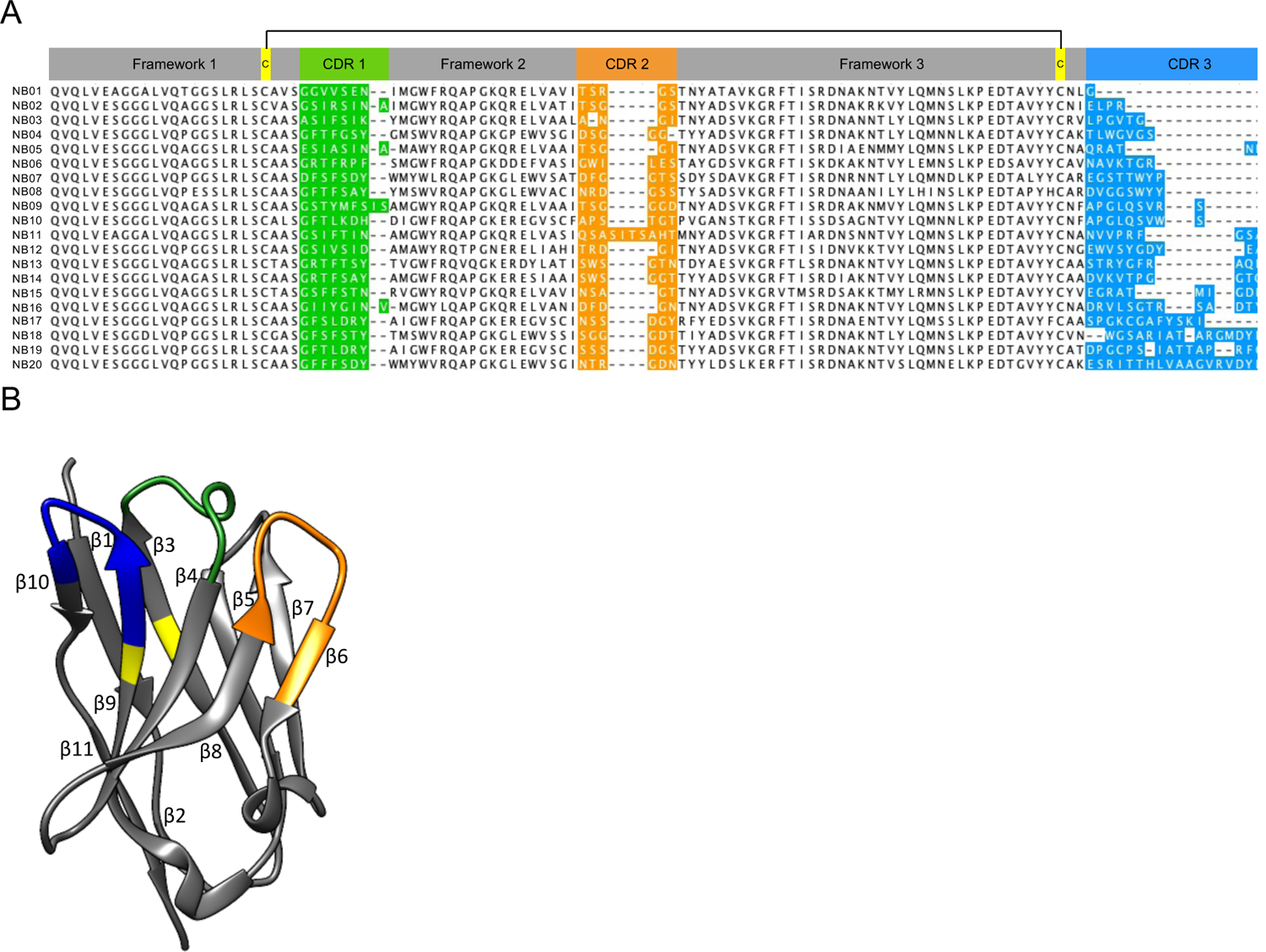
Nanobody families and overall structure. (A) Sequence alignment of the 20 selected antibodies. Six large nanobody families dominated the phage display output: over 60% of the nanobodies belong to these families. Nanobodies within the same family are identified by their shared font colour. Sixteen unique families consisted of only a few nanobody clones. The complementarity determining regions (CDR), CDR1, CDR2 and CDR3 are aligned. (B) Nanobody structure. The anti-green fluorescent protein (GFP) nanobody structure (PDB: 3OGO) is used to exemplify the nanobody fold. The three CDR1-3 loops are coloured green, orange and blue respectively and the location of disulphide bonds in yellow. (**Associated with Figure S1**).

### Biophysical analysis of p16 nanobodies

We carried out a range of orthogonal biophysical techniques to confirm the structural integrity of the selected nanobodies. First, far-UV circular dichroism (far-UV CD) was used to assess nanobody secondary structure (**Figure 2A, D** and **Figure S2A**). All exhibited characteristic minima at 218 nm corresponding to the antiparallel beta sheet forming the core immunoglobulin fold. Differences in CD signal were most likely caused by the different structures adopted by the CDR loops, which differ in length and sequence, between the nanobody families.

**Figure 2.**
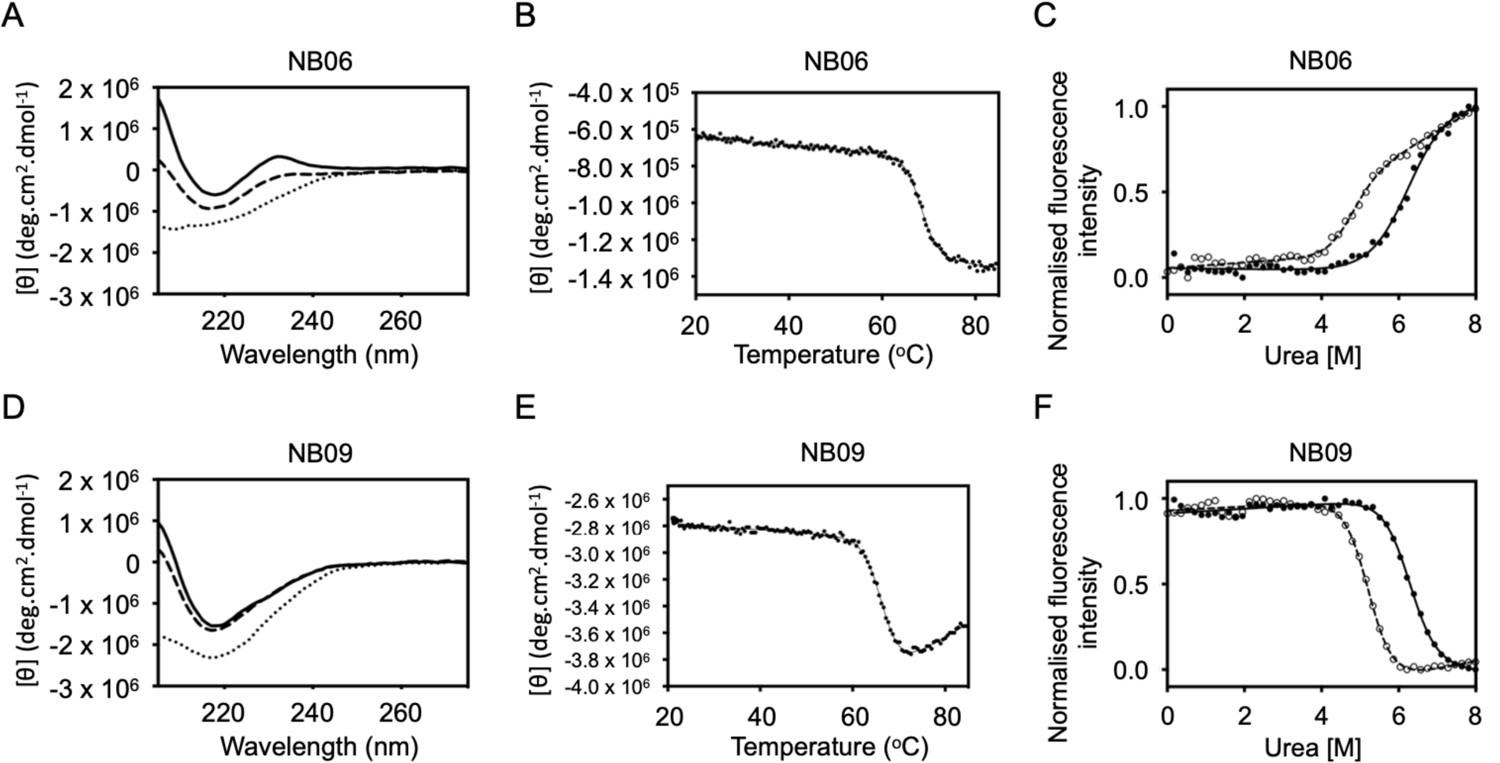
Nanobody characterisation. (A) Far-UV circular dichroism (far-UV CD) spectra, (B) thermal stability, (C) Chemical denaturation monitored by tryptophan fluorescence. Of the 19 nanobodies tested, 15 nanobodies showed a cooperative two-state unfolding. (**Associated with Figure S2 and Table S1**).

**Table 1.** Thermodynamic stability of p16 mutants in the presence of either NB06 or NB09. **Related to Figure 4**.

| Construct | Nanobody | $D_{50\%}^1$ | $m^1$ | $\Delta G_{unf}^2$ |
| --- | --- | --- | --- | --- |
|  |  | (M) | (kcal mol<sup>-1</sup> M<sup>-1</sup>) | (kcal mol<sup>-1</sup>) |
| **R24P** |  | <sup>-3</sup> | - | - |
| **R24P** | NB06 | 1.03 ± 0.05 | <sup>-4</sup> | 1.71 ± 0.09 |
| **R24P** | NB09 | 1.43 ± 0.07 | <sup>-4</sup> | 2.37 ± 0.12 |
| **Q50R** |  |  |  |  |
| **Q50R** | NB06 | 1.20 ± 0.05 | <sup>-4</sup> | 1.99 ± 0.07 |
| **Q50R** | NB09 | 1.05 ± 0.06 | <sup>-4</sup> | 1.74 ± 0.10 |
| **D74N** |  | 1.03 ± 0.05 | <sup>-4</sup> | 1.71 ± 0.09 |
| **D74N** | NB06 | 2.14 ± 0.01 | 1.69 ± 0.05 | 3.55 ± 0.05 |
| **D74N** | NB09 | 2.22 ± 0.01 | 1.93 ± 0.06 | 3.69 ± 0.05 |
| **P81L** |  | 1.77 ± 0.01 | 2.06 ± 0.02 | 2.94 ± 0.04 |
| **P81L** | NB06 | 1.80 ± 0.01 | 1.84 ± 0.05 | 2.99 ± 0.04 |
| **P81L** | NB09 | 1.75 ± 0.01 | 2.15 ± 0.05 | 2.91 ± 0.04 |
| **D84N** |  | 1.79 ± 0.01 | 2.10 ± 0.02 | 2.97 ± 0.04 |
| **D84N** | NB06 | 2.49 ± 0.01 | 1.77 ± 0.03 | 4.13 ± 0.05 |
| **D84N** | NB09 | 2.61 ± 0.01 | 2.00 ± 0.06 | 4.33 ± 0.05 |
| **H98P** |  | <sup>-3</sup> | - | - |
| **H98P** | NB06 | <sup>-3</sup> | - | - |
| **H98P** | NB09 | <sup>-3</sup> | - | - |
| **P114L** |  | 1.04 ± 0.06 | <sup>-4</sup> | 1.72 ± 0.10 |
| **P114L** | NB06 | 0.69 ± 0.33 | <sup>-4</sup> | 0.11 ± 0.55 |
| **P114L** | NB09 | 1.38 ± 0.07 | <sup>-4</sup> | 2.29 ± 0.12 |
| **V126D** |  | <sup>-3</sup> | - | - |
| **V126D** | NB06 | <sup>-3</sup> | - | - |
| **V126D** | NB09 | <sup>-3</sup> | - | - |

<sup>1</sup> Denaturation experiments were performed at 25 °C in 50 mM Tris-HCl pH 7.5, 100 mM NaCl. The concentration of each protein was 2 μM. Data were fitted to a two-state model to obtain the midpoint of unfolding, D50%, and the $m$ -value. N = 3 ± SD.
<sup>2</sup> The free energy of unfolding, $\Delta G_{unf}$ , was calculated as the product of the midpoint of unfolding, D50%, and the wild-type $m$ -value (1.66 ± 0.02). For those datasets (R24P, Q50R, D74N and P114L) that lack a significant pre-transition baseline, the denaturation curves were fitted by fixing $m$ to the wild-type value.
<sup>3</sup> Not determined due to the absence of a clear unfolding transition.
<sup>4</sup> Due to the very low stability resulting in an absence of a pre-transition baseline, the denaturation curves were fitted by fixing $m$ to the wild-type value.

Next, thermal stability was measured by monitoring the ellipticity at 218 nm that corresponds to the signal from the anti-parallel beta-sheet of the immunoglobulin fold as a function of temperature (**Figure 2B, E** and **Figure S2B**). The nanobodies all exhibited a single unfolding transition. The thermal stability for the majority of the nanobodies was high, with midpoints of unfolding between 60 °C and 80 °C (**Table S1**). There was no correlation between the length of CDR3 and stability, suggesting that differences in CDR3 sequence were not the only factor in determining nanobody stability. The ability of the nanobodies to refold was also investigated. A far UV-spectrum was recorded after denaturation at 80 °C, the nanobody was allowed to cool to 20 °C and a second spectrum was recorded. Although there was a slight loss of signal for some nanobodies, NB04, NB05, NB12 and NB15 recovered all their signal. Interestingly NB07 also regained the full signal, suggesting that despite the low T_m_ this nanobody was able to refold. Some nanobodies (E.g. NB06 and NB13) showed a marked loss in signal upon cooling, which may be due to irreversible unfolding of the immunoglobulin scaffold and loss of the core disulphide bond.

Lastly, to investigate the chemical stability of the nanobodies, tryptophan fluorescence was monitored as a function of urea concentrations. The core disulphide bond increases stability, such that to assess intracellular nanobody activity it is critical to determine whether the loss of this core disulphide bond is detrimental to nanobody stability. Of the 19 nanobodies tested, 15 showed a cooperative two-state unfolding (**Figure 2C, F** and **Figure S2C** and **Table S1**). The stabilities of the nanobodies are high, with many having a midpoint of denaturation (D50%) of 5-7 M urea. There were modest reductions in stability of 1-2 M due to the loss of the core disulphide bond, but all nanobodies remained folded up to around 4 M urea. From these analyses the six nanobodies (NB03, NB05, NB06, NB09, NB16 and NB17) that had the highest thermodynamic stabilities, and fluorescence profiles that gave a flat pre-transition baseline, were selected for characterisation of their effects on p16.

### Nanobodies stabilize wild-type and mutant p16

p16 is fully unfolded at approximately 3M urea, a urea concentration at which the six selected nanobodies will remain folded. Wild-type p16 exhibited a two-state urea-induced unfolding profile when monitored by tryptophan fluorescence, (**Figure 3**). The D50% was 1.94 ± 0.01 M urea and the *m*-value (a parameter that is a measure of the change in solvent-accessible surface area upon unfolding) was 1.66 ± 0.02 kcal mol^−1^ M^−1^. The D50% and *m*-value did not change significantly in the presence of four of the nanobodies NB03, NB05, NB16 or NB17 (**Figure 3** and **Table S2**). The midpoint of unfolding of p16 greatly increased in the presence of nanobodies NB06 and NB09 (by almost 45 % to 2.80 ± 0.01 M urea and by over 60 % to 3.11 ± 0.04 M urea respectively). The free energy of unfolding increased from 3.22 ± 0.04 kcal mol^−1^ for wild-type p16 to 4.65 ± 0.06 kcal mol^−1^ and 5.16 ± 0.09 kcal mol^−1^ when bound to NB06 and NB09, respectively. Given their superior ability to stabilize wild type p16, NB06 and NB09 were taken forward to assess their abilities to stabilize clinically relevant mutant forms of p16.

**Figure 3.**
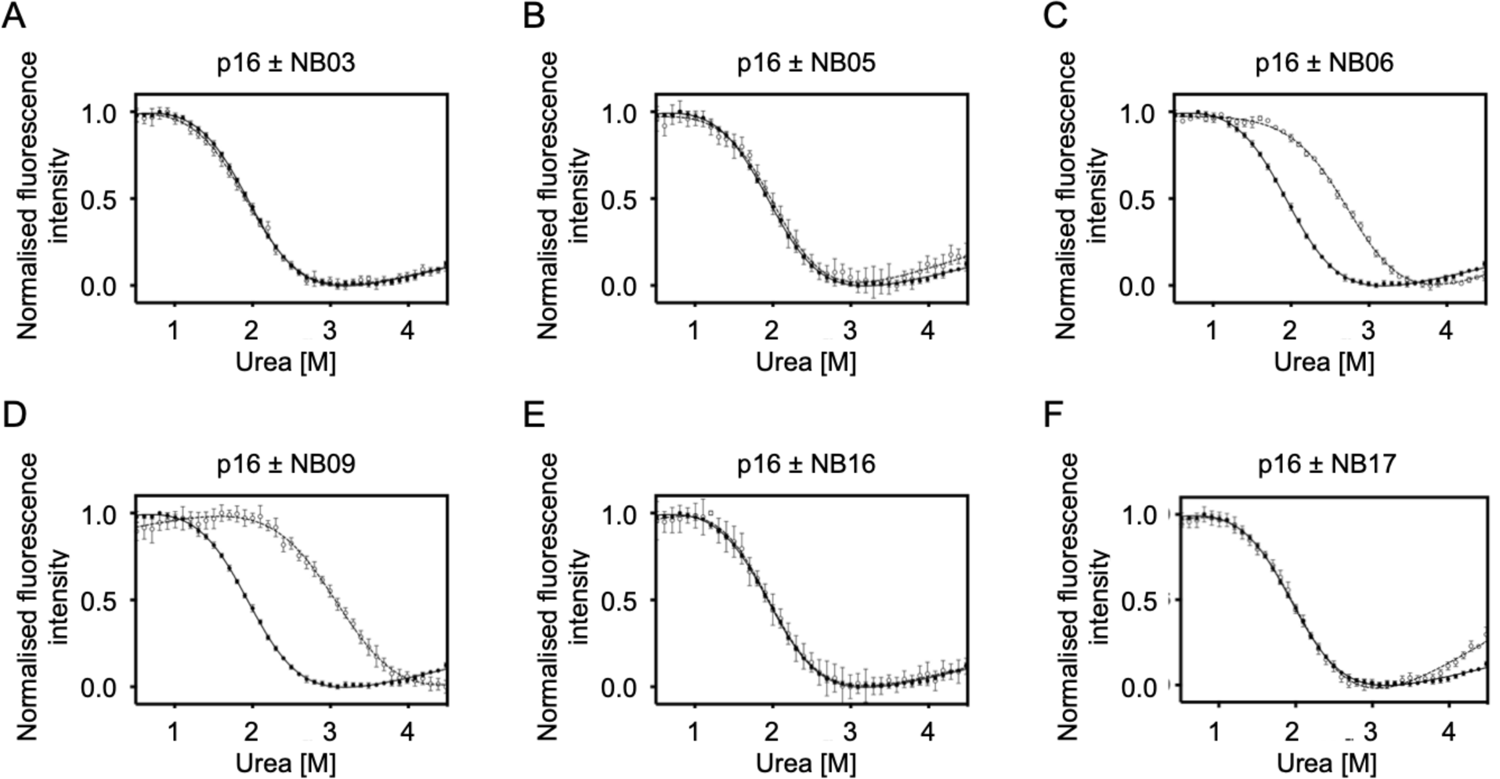
Nanobodies stabilize wild-type and mutant p16. Fluorescence-monitored denaturation curves of wild-type p16 alone and in complex with six different nanobodies. Chemical denaturation was performed at 25 °C in 50 mM Tris-HCl buffer pH 7.5, 100 mM NaCl. Excitation wavelength was 296 ± 10 nm and emission wavelength 360 ± 20 nm. Experiments were performed in triplicate and the data fitted to a two-state model (Prism). Filled circles, without dithiothreitol (DTT); open circles, buffer supplemented with 1 mM DTT.

To assess the ability of NB06 and NB09 to stabilize mutant p16 proteins, a set of eight cancer-associated p16 mutations were selected based on their dispersed locations throughout the p16 structure and different effects on p16 stability (**Figure 4A** and **Table 1**) (Tate et al., 2019). To determine denaturation profiles, mutant and wild-type p16 were first unfolded in 4 M urea and then refolded by aliquoting into a range of urea concentrations. As has been observed previously (Boice and Fairman, 1996; Byeon et al., 1998; Tevelev et al., 1996; Yang et al., 1995; Yuan et al., 1999; Zhang and Peng, 1996), some but not all cancer-associated mutants are capable of folding to the native structure (**Figure 4B** and **Table 1**). The urea denaturation profiles of mutants p16^D74N^, p16^P81L^ and p16^D84N^ were similar to that of wild-type p16, indicating that they can refold to the native structure, though they have somewhat reduced stabilities. p16^D74N^ and p16^D84N^ are at the CDK-binding interface and therefore were not expected to drastically compromise the fold (**Figure 4A**). In contrast, p16^R24P^ and p16^H98P^ (both solvent-exposed) and p16^V126D^ (buried) showed little or no change in fluorescence with increasing urea concentration, indicating that they did not refold. For p16^Q50R^ (solvent-exposed) and p16^P114L^ (both buried), some aggregation was observed below 1 M urea and the unfolding profiles were relatively noisy but could nevertheless be fitted to a two-state equation which gave low midpoints of unfolding.

**Figure 4.**
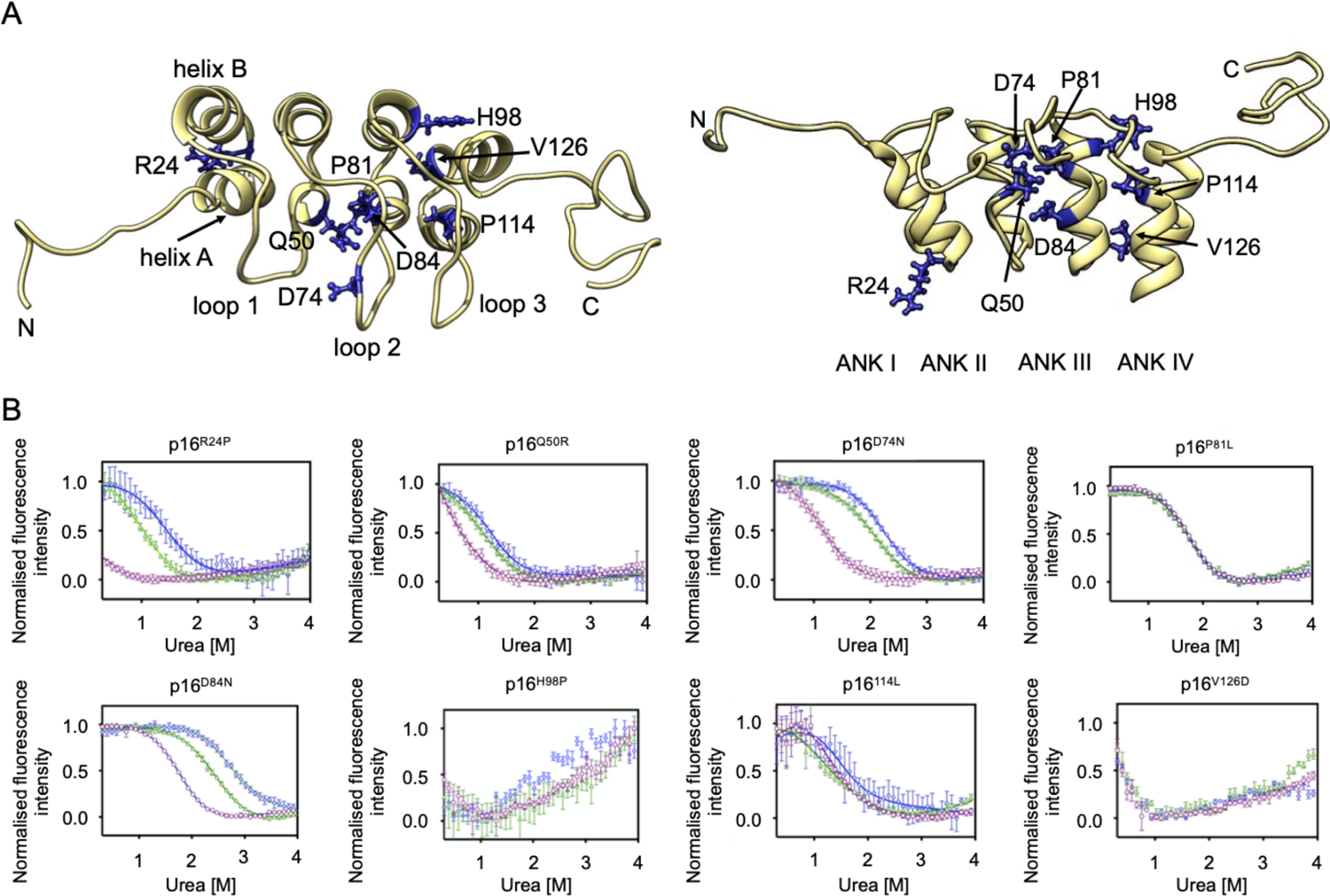
p16 structure and characterisation of cancer-associated mutations. (A) Structure of p16 (PDB: 2A5E) showing location of cancer-associated mutants. For clarity, the lower panel is rotated 90° with respect to the upper panel. The two helices within the first ankyrin repeat are labelled A and B. (B) Fluorescence-monitored denaturation curves of p16 mutants alone and in complex with nanobodies NB06 and NB09. Data was normalised to maximum and minimum fluorescence intensity. Chemical denaturation was performed at 25 °C in 50 mM Tris-HCl buffer pH 7.5, 100 mM NaCl. Excitation wavelength was 295 ± 10 nm and emission wavelength 360 ± 20 nm. Purple open circles, p16 mutant alone; green open triangles, p16 mutant and NB06; blue open diamonds, p16 mutant and NB09. Experiments were performed in triplicate and the data fitted to a two-state model (Prism). (**Associated with Figure S3**).

Next, we characterized the interactions of NB06 and NB09 with the eight p16 cancer-associated variants (**Table 1, Figure S3**). Both nanobodies stabilize the p16 variants p16^R24P^, p16^Q50R^, p16^D74N^ and p16^D84N^. These sites are all solvent exposed, R24 and D74 being in loops, Q50 and D84 within helices. The stabilising effect was particularly dramatic for p16^R24P^, which showed no cooperative unfolding in the absence of nanobodies but displayed a cooperative unfolding behaviour in the presence of either nanobody. NB09 generally increased the stability to a greater extent than did NB06; this behaviour is like that of wild type p16, which also showed greater stability bound to NB09 than to NB06 (**Table S2**). In contrast to these four variants, the stability of p16^P81L^, a buried residue that precedes helix A of ANKIII, was strikingly unaffected by either nanobody. For the three remaining variants tested, p16^H98P^ (at the end of helix B in ANKIII), p16^P114L^ (at the start of helix A in ANKIV), and p16^V126D^ (at the start of helix B in ANKIV), all of which showed highly compromised folding, there was no detectable effect of either nanobody. It can be hypothesized that the structures of these p16 mutants are too compromised for the nanobodies to be able to rescue them.

In summary our analysis shows that NB06 and NB09 stabilize wild-type p16 and distinguish different mutant p16 proteins. Mutants p16^R24P^, p16^Q50R^, p16^D74N^ and p16^D84N^ were stabilized by the nanobodies, whereas p16^P81L^, p16^H98P^, p16^P114L^ and p16^V126D^ were not. Interestingly, two of the former set, p16^R24P^ and p16^Q50R^, are highly unstable and misfolded mutant proteins, indicating that rescue is not limited to only mildly destabilizing mutations.

### Nanobody binding to p16 is compatible with formation of a ternary nanobody-p16-CDK complex

We next used surface plasmon resonance (SPR) and isothermal titration calorimetry (ITC) to compare the binding of NB06 and NB09 to p16. In the SPR experiments, GSTp16 was captured as a ligand on the CM5 chip with coupled anti-GST antibody, and untagged nanobodies NB06 and NB09 were flowed over as an analyte. This analysis revealed that NB09 binds slightly tighter to p16 than NB06 with measured K_d_ values of 7.52 ± 1.8 nM and 19.98 ± 6.1 nM, respectively. Dissection of the kinetics of the interactions revealed that NB06 has a 10-fold higher rate of dissociation from p16 (k_off_ = 4.2e-02 s^-1^) than NB09 (k_off_ = 3.30e-03 s^-1^) but their association rates (k_on_) are comparable, calculated as 1.98e+06 M^-1^s^-1^ (NB06) and 3.40e+06 M^-1^s^-1^ (NB09) respectively (**Figure 5A**). Comparable K_d_ values were determined from the equilibrium binding measurements (24.7 ± 9.6 nM and 10.2 ± 1.8 nM for binding of NB06 and NB09 respectively). ITC confirmed the 1:1 stoichiometry of the nanobody-p16 interaction and their nanomolar potency. NB09 displayed slightly higher affinity for p16 (8.0 ± 0.37 nM) than NB06 (26 ± 5.2 nM) (**Figure 5B**). Notably, the NB06-p16 interaction was endothermic (Δ*H* = +1.7 ± 0.02 kcal mol^-1^), whereas the NB09-p16 interaction was exothermic (Δ*H* = −9.0 ± 0.03 kcal mol^-1^).

**Figure 5.**
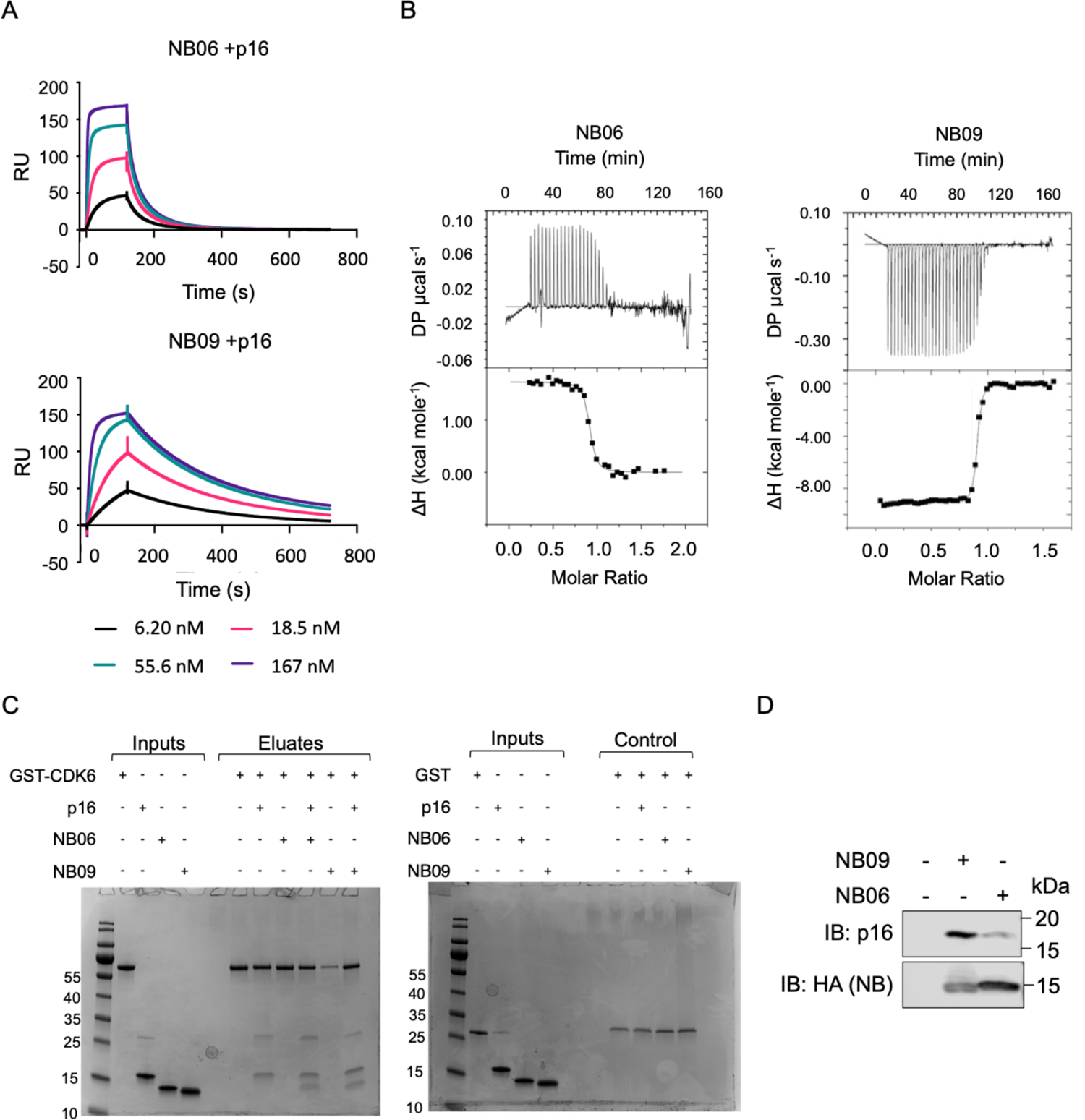
Nanobody and CDK binding to p16 are compatible. p16 binding to NB06 and NB09 as measured by surface plasmon resonance (A) and isothermal titration calorimetry (ITC) (B). (A) GSTp16 was immobilized on the chip via anti-GST antibody coupling and the NB06 and NB09 analytes were assayed in triplicate over a 5-point serial dilution. Dissociation and kinetic rate constants were derived by using the Biacore S200 Evaluation Software. (B) ITC titration of NB06 (180 μM in syringe, LHS panel) and NB09 (128 μM, in syringe, RHS panel) into p16. p16 was at a concentration in the cell of 19 μM or 18 μM respectively. Data was fitted to one-site binding model. ITC experiments were carried out with one biological replicate and values are quoted ± SD. (C) Pulldowns using recombinant proteins. GSTCDK6 was co-incubated with p16 or/and with p16 in the presence of a nanobody NB06 or NB09 and then analyzed by SDS-PAGE and subsequent InstantBlue staining. GST was used as a control. (D) Nanobodies bind endogenous p16 in HEK293T cells. HEK293T cells were transfected with either NB09 or NB06 and resultant cell lysates immunoprecipitated with anti-HA beads prior to analysis by western blot. (**Associated with Figure S4** (uncropped gel)).

The nanobodies were selected from screens against p16 and p16-CDK complexes such that they should not preclude p16-CDK association. Bead-bound recombinant GST-CDK6 was incubated with p16 in the presence or absence of NB06 or NB09 and bead-bound fractions were analysed by SDS-PAGE (**Figure 5C**). Under those conditions GST-CDK6 forms a ternary complex with p16 and either nanobody. CDK6 does not associate with either nanobody in the absence of p16. Analytical size exclusion chromatography also demonstrated formation of a stable ternary complex (**Figure S4A**).

We used Homogenous Time Resolved Fluorescence (HTRF) to assess whether binding of NB06 or NB09 impacts the affinity of the interaction between p16 and CDK4 or CDK6. CDK4 and CDK6 bind to p16 with K_d_ values in the low nanomolar range and within the experimental parameters nanobody addition to the incubation did not perturb the interaction (**Figure S4B**). However, because these dissociation constants correspond to less than the concentration of bait (GST-CDK) used in the HTRF assay, they represent upper estimates for the affinity of each interaction.

Lastly, we investigated the interactions between nanobodies and p16 in a cellular setting. We assessed the interactions between nanobodies and endogenous (wild-type) p16 in HEK293T cells (**Figure 5D, Figure S4C**). NB09 pulled down more p16 than did NB06 despite the apparent lower expression of NB09 relative to NB06. When corrected for expression levels, NB09 pulled down *circa* 8-fold more p16 than did NB06 (**Figure 5D**). The larger effect on p16 levels of NB09 relative to NB06 is consistent with the greater stabilization of purified p16 protein by NB09 compared with NB06.

### Crystal structure of p16 in complex with nanobody NB09 rationalizes mutant p16 behavior

The crystal structure of NB09 in complex with p16 was determined to 1.79 Å resolution (**Figure 6A, Table S3**). Within the crystal lattice the nanobody forms a large proportion of the key contacts with a major interface forming between adjacent nanobody framework regions (**Figure S5**). The crystal packing is similar to that seen in the crystal structure of the flexible MazE protein in complex with a nanobody (Loris et al., 2003).

**Figure 6.**
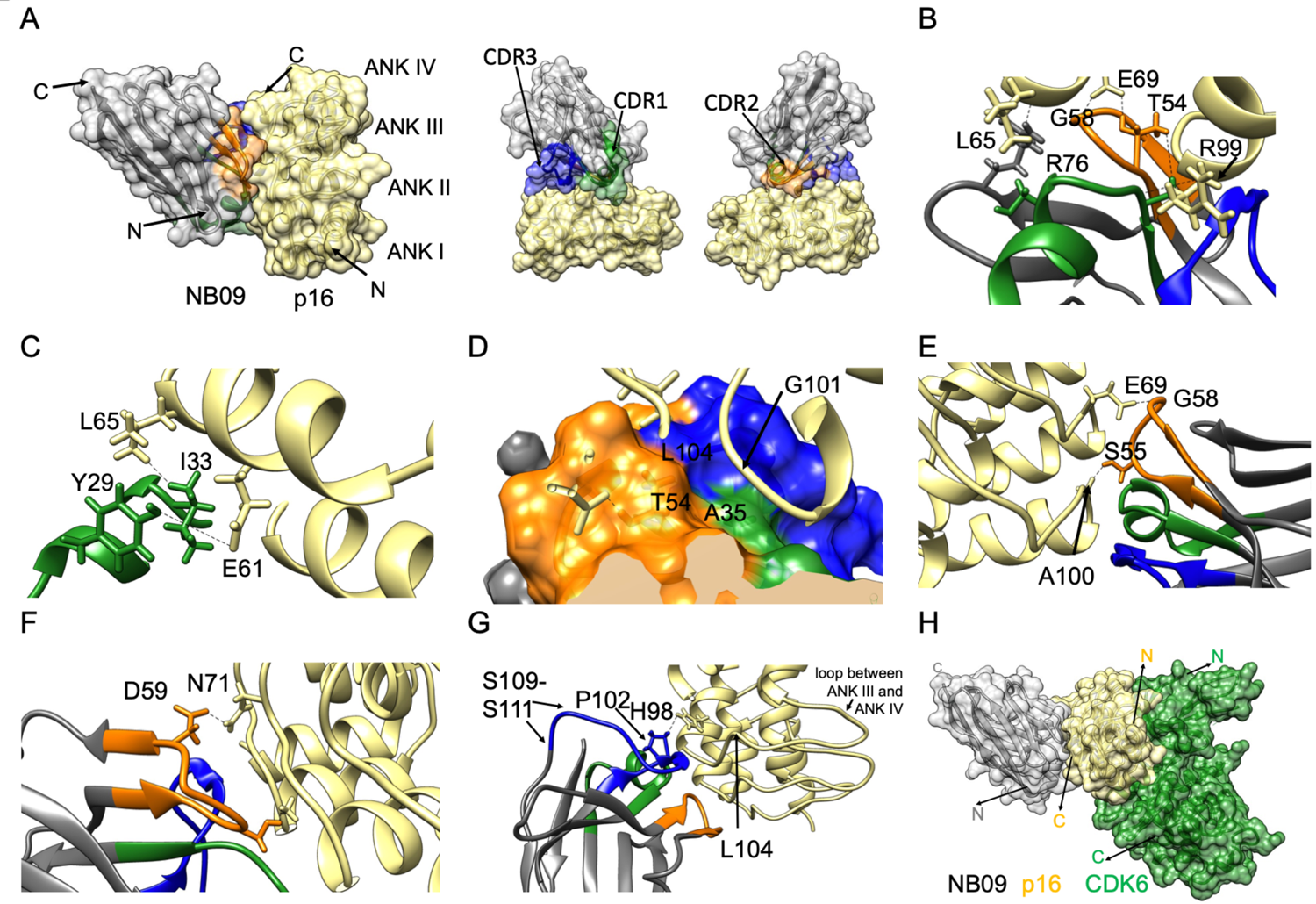
Crystal structure of the p16-NB09 nanobody complex. (A) The crystal structure of NB09 (grey) in complex with p16 (straw). The structures in the right hand side panel are related by a rotation of 180° around a vertical axis. (B-G) Close up views of the NB09-p16 interface. Selected residues are labelled. The nanobody CDR1, CDR2 and CDR3 sequences are highlighted in green, orange and blue, respectively. (H) CDK6-p16-NB09 model. NB09 in complex with p16 is superimposed onto the CDK6 crystal structure extracted from the structure of the p16-CDK6 complex (PDB: 1BI7). (**Associated with** Figures S5 and S6).

The N- and C-terminal regions of p16 are predicted to be unstructured and, despite the improved resolution when compared to the previously determined p16-CDK6 complex (PDB: 1BI7 at 3.4 Å resolution (Russo et al., 1998)), the N-(M1-M9) and C-(G135-D156) terminal residues of p16 remain unresolved in the p16-NB09 structure. However, the nanobody structure is well resolved and the chain can be traced from residues Q1-S122.

Analysis of the complex reveals a free solvent-accessible surface area (Mitternacht, 2016) of the nanobody paratope of 684.7 Å^2^, similar to the average for other published nanobody complexes (768.5 ± 201.0 Å^2^) (Mitchell & Colwell, 2018). Furthermore, shape complementary analysis (Lawrence & Colman, 1993) of both the nanobody paratope and p16 epitope was 0.727, consistent with the average for other published nanobody structures (0.72 ± 0.07) (Mitchell & Colwell, 2018).

In the p16-NB09 structure, residues within each CDR loop contact p16: from the C-terminal end of CDR1 (green) throughout CDR2 (orange) and towards the N-terminal end of CDR3 (blue) (**Figure 6A, B)**. The NB09 CDR1 is unusually long (**Figure 1A**, (Mitchell and Colwell, 2018)) and forms an alpha helix with two turns that is orientated perpendicular to ANK2 and ANK3 and surface exposed. Following the helix I33 packs with p16 L65 (towards the C-terminal end of ANK2). In the vicinity the sidechain of NB09 Y29 points towards p16 E61 (**Figure 6C**). A35 at the start of framework 2 together with T54 (from CDR2) and L104 (from CDR3) form a pocket into which p16 G101 (just following the end of ANK3) puckers (**Figure 6D**). These residues also engage in hydrogen bonds, between the backbone amide of NB09 A35 and the backbone carbonyl of p16 R99, the sidechain hydroxyl of T54 and the sidechain of p16 E69, and between the sidechain of R76 (within framework 3) and the backbone carbonyl of p16 L65 (**Figure 6B**).

The CDR2 loop has an unusual conformation permitted by the glycine-rich SGGG sequence that enables the amide moieties of S55 and G58 to form hydrogen bonds with respectively the backbone carbonyl of p16 A100 and the sidechain of E69 (**Figure 6E**). A water coordinates a network of hydrogen bonds between the side chain of NB09 S55 and the carbonyl oxygens of L64 and A68 respectively at the C-terminal end of helix 2 of ANK2 and the beginning of the loop between ANK 2 and ANK 3 of p16. At the end of CDR2, D59 makes a direct interaction with the side chain of p16 N71 and via a water across to the sidechain of p16 E69 (**Figure 6F**).

In a number of nanobody-protein complexes, the CDR3 loop is responsible for a large proportion of the binding contacts at the nanobody:protein interface (Desmyter et al., 1996). Additional interactions made by CDR3 also contribute to the interface. P102 and G103 found at the N-terminal end of the CDR3 loop form a tight turn which places the loop onto a shallow pocket on the surface of p16 facing p16 H98 (**Figure 6G**). The carbonyl oxygen of G103 also makes a water-mediated interaction across to the backbone nitrogen of L104 in the p16 ANK3 to ANK4 loop. Residues S109-S111 at the C-terminal end of CDR3 are flexible.

As described above, mutants p16^R24P^, p16^Q50R^, p16^D74N^ and p16^D84N^ are stabilized by NB09. The structure of the NB09-p16 complex rationalises the ability of NB09 to stabilise this subset of p16 mutants. p16 R24, Q50, D74 and D84 are all solvent exposed and are distant from the NB09-p16 interface. Interactions between NB09 and p16 are able to assist in stabilising these p16 mutant structures because these mutations neither impact the integrity of the core of the p16 fold nor alter the residues at the nanobody-p16 interface. As revealed by the structure of the CDK6-p16 complex (Russo et al., 1998), mutations to p16 R24, D74 and D84 impact CDK6 activity because they are at the p16-CDK6 interface. Q50 is located in the vicinity of D84.

Four p16 mutants, p16^P81L^, p16^H98P^, p16^P114L^ and p16^V126D^ were not stabilised by either nanobody. P81, P114 and V126 are buried, and H98 is surface exposed within helix 2 of ANK3 at the CDK6 interface. Given their locations and the amino acid changes, all these mutations would be expected to destabilise the p16 fold, and indeed that is what is observed. It is hypothesized that for these mutants (and in contrast to p16 mutants p16^R24P^ and p16^Q50R^, that are also highly unstable and misfolded mutant proteins) nanobody binding is unable to rescue p16 structural integrity.

The epitope recognised by NB09 sits opposite the flexible loops that link adjacent ankyrin repeats of p16 with the majority of the binding interface being situated at the interfaces between ANK2 and ANK3 and between ANK3 and ANK4. Thus, the nanobody binding site on p16 is on the opposite face to where it binds CDK6 (**Figure 6H**). An alignment of the two p16 crystal structures reveals minor differences between them (RMSD of 0.693 Å) indicating that binding of NB09 to p16 does not significantly alter the overall p16 fold (**Figure S6**). As NB09 does not block the CDK6 binding site nor change the overall structure of p16, the model confirms our experimental results that p16 binds to CDK6 in the presence of NB09. Taken together, we propose that p16 bound to NB09 is still capable of inhibiting CDK6 (and by extension CDK4). Thus, NB09 meets the criteria to be used as a potential pharmacological chaperone.

## Significance

Camelid single-domain antibodies (nanobodies) possess a range of properties that make them useful tools in structural biology, and several are under development as therapeutics in diseases associated with protein misfolding. Here we describe the identification and characterisation of nanobodies to stabilize cancer-associated mutations in the tumor suppressor protein p16. p16 plays a critical role in regulation of the cell cycle by binding and inhibiting cyclin-dependent kinases 4 and 6 (CDK4 and CDK6), thereby blocking pRb phosphorylation and entry into S phase. p16 is a thermodynamically unstable protein, and hundreds of cancer-associated missense mutations have been identified, located throughout p16, the majority of which cause loss of function by destabilising the protein’s structure. We show that the nanobodies bind to p16 with nanomolar affinities, that binding results in a large increase in the thermodynamic stability of p16, and that the stabilisation of p16 is also achieved in the cellular context. Furthermore, the nanobodies were also able to stabilize a range of different cancer-associated p16 mutations located at sites throughout the structure. We co-crystallised one of the nanobodies in complex with p16. The structure revealed that the nanobody binds to the opposite face of p16 to the CDK-binding interface and the CDK-binding ability of nanobody-bound p16 was subsequently confirmed by solution studies. These findings demonstrate that nanobodies can bind and stabilize conformationally flexible proteins as well as restore the stability of destabilising cancer-associated mutations, pointing to their potential use as pharmacological chaperones to rescue p16 function in the cell.

## Acknowledgements

X-ray crystallography was carried out with the support of Diamond Light Source on beamlines I03 (allocation MX13587). We thank E. Pardon and J. Steyaert (Structural Biology Research Center, Brussels) and INSTRUCT, part of the European Strategy Forum on Research Infrastructures (ESFRI) and the Hercules Foundation Flanders for their support with the Nanobody discovery and R. Owens for plasmid deposition in Addgene. We thank S. Hallett for assistance with preparation and initial construct characterization, A Baslé (University of Newcastle) for support with data collection and processing, E. De Genst (University of Cambridge, Department of Chemistry) for initial advice on nanobody production, and H. Laman (University of Cambridge, Department of Pathology). This research was supported by the MRC (Grant MR/N009738/1, MWP). ODB and SH were supported by CRUK Cambridge Cancer Centre PhD studentships. LSI and GZ acknowledge the support of a fellowship from the UK Medical Research Foundation (Grant C0385) and a project grant from the Pancreatic Cancer Research Foundation. ODB acknowledges the support of a Fieldwork Fund grant from University of Cambridge School of the Biological Sciences.

## Author Contributions

Owen Burbidge: Conceptualization, methodology, investigation (cloning, construct design, protein production, phage display, biophysical analysis (SEC, CD, chemical and thermal denaturation crystallography)), writing original draft, review and editing.

Martyna Pastok: Conceptualization, investigation (cloning, construct design, protein production, SPR, HTRF and immunoprecipitation experiments, X-ray crystallography) and writing original draft, review and editing.

Grasilda Zenkevičiūtė: Investigation and writing (protein production, ITC and cell-based experiments)

Samantha Hodder: Investigation and writing (protein production, ITC and cell-based experiments)

Martin Noble: Conceptualization, resources, supervision, and funding acquisition. Jane Endicott: Conceptualization, resources, supervision, and funding acquisition, review and editing.

Laura Itzhaki: Conceptualization, resources, supervision, funding acquisition, writing original draft, review and editing.

## Materials and Methods

### Key resources table

| REAGENT or RESOURCE | SOURCE | IDENTIFIER |
| --- | --- | --- |
| **Antibodies** |  |  |
| Tb-anti-GST antibody | CisBio | 61GSTTLB |
| Anti-GST antibody (SPR analysis) | GST Capture Kit GE Lifescience | BR100223 |
| Streptavidin-Alkaline Phosphatase | Promega | V5591 |
| CaptureSelect Biotin anti-C-Tag conjugate | Invitrogen | 7103252100 |
| Anti-CDKN2A/p16INK4a | Abcam | Ab108349 |
| HA-Tag (C29F4) | Cell Signalling Technology | 3724 |
| Polyclonal swine anti-rabbit immunoglobulins/HRP | Dako | P0399 |
| Polyclonal goat anti-mouse immunoglobulins/HRP | Dako | P0447 |
| **Bacterial and virus strains** |  |  |
| *E. coli* DH5α | Invitrogen | 18265-017 |
| *E. coli* BL21STAR™ (DE3) | ThermoFisher Scientific | C601003 |
| *E. coli* Rosetta2 (DE3) pLYS-S | Novagen | 709564 |
| *E. coli* DH10EmBacY *E. coli* | Geneva Biotech | Supplied as part of MultiBac™ Expression System Kit |
| *E. coli* TG1 | Lucigen | 60502-1 |
| *E. coli* OmniMaxII | ThermoFisher Scientific | C854003 |
| *E. coli* Lemo21 (DE3) | New England Biolabs | C2528J |
| VCSM13 helper phage | Stratagene (Agilent) | 200251 |
| **Biological samples** |  |  |
| N/A |  |  |
| **Chemicals, peptides, and recombinant proteins** |  |  |
| Streptavidin-labelled XL665 | CisBio | 610SAXLB |
| **Critical commercial assays** |  |  |
| MultiBac™ Expression System Kit | Geneva Biotech | N/A |
| GST Capture Kit | GE Lifescience | BR100223 |
| Morpheus crystallization screen | Molecular Dimensions | MD1-123 |
| GeneJuice® Transfection reagent | Merck Millipore | 70967 |
| Insect-Xpress insect cells medium | Lonza | BELN12-730Q |
| **Deposited data** |  |  |
| Crystal structure of p16-NB09 nanobody complex | This paper | PDB entry 7OZT |
| **Experimental models: Cell lines** |  |  |
| *Spodoptera frugiperda* Sf9 cells | OET | 600100 |
| **Experimental models: Organisms/strains** |  |  |
| **Oligonucleotides** |  |  |
| See Table S3 for oligonucleotide sequences |  |  |
| **Recombinant DNA** |  |  |
| pACEBac1-GST-CDK4 | (Hallett *et al.*, 2017) |  |
| pACEBac1-GST-CDK6 | (Hallett *et al.*, 2017) |  |
| pGEX6P1-GST-p16 | (Hallett *et al.*, 2017) |  |
| pMESy4 |  | Genbank KF415192 |
| pMESy4-NB01 | This paper |  |
| pMESy4-NB02 | This paper |  |
| pMESy4-NB03 | This paper |  |
| pMESy4-NB04 | This paper |  |
| pMESy4-NB05 | This paper |  |
| pMESy4-NB06 | This paper |  |
| pMESy4-NB07 | This paper |  |
| pMESy4-NB08 | This paper |  |
| pMESy4-NB09 | This paper |  |
| pMESy4-NB10 | This paper |  |
| pMESy4-NB11 | This paper |  |
| pMESy4-NB12 | This paper |  |
| pMESy4-NB13 | This paper |  |
| pMESy4-NB14 | This paper |  |
| pMESy4-NB15 | This paper |  |
| pMESy4-NB16 | This paper |  |
| pMESy4-NB17 | This paper |  |
| pMESy4-NB18 | This paper |  |
| pMESy4-NB19 | This paper |  |
| pMESy4-NB20 | This paper |  |
| pOPINE | (Berrow *et al.*, 2007) | Addgene 26043 |
| pOPINF | (Berrow *et al.*, 2007) | Addgene 26042 |
| pOPINF-p16 | This paper |  |
| pOPINF-p16<sup>L37S/L121R</sup> | This paper |  |
| pOPINF-p16<sup>NTA</sup> | This paper |  |
| pOPINF-p16<sup>CTA</sup> | This paper |  |
| pOPINF-HA_NB09 | This paper |  |
| pOPINF-HA_NB06 | This paper |  |
| pOPINJ | (Berrow *et al.*, 2007) | Addgene 26045 |
| pOPINJ-p16 | This paper |  |
| pOPINM | (Berrow *et al.*, 2007) | Addgene 26044 |
| pOPINM-p16 | This paper |  |
| pOPINS3C | (Bird, 2011) | Addgene 41115 |
| POPINS3C-p16 | This paper |  |
| pMA | Life technologies – GeneArt cloning vector |  |
| pRSET A | ThermoFisher | V35120 |
| pRSET A-p16<sup>R24P</sup> | This paper |  |
| pRSET A-p16<sup>Q50R</sup> | This paper |  |
| pRSET A-p16<sup>D74N</sup> | This paper |  |
| pRSET A-p16<sup>P81L</sup> | This paper |  |
| pRSET A-p16<sup>D84N</sup> | This paper |  |
| pRSET A-p16<sup>H98P</sup> | This paper |  |
| pRSET A-p16<sup>P114L</sup> | This paper |  |
| pRSET A-p16<sup>V126D</sup> | This paper |  |
| **Software and algorithms** |  |  |
| Biacore S200 Evaluation Software 1.0 | GEHealthcare | N/A |
| GraphPad Prism 9 | GraphPad | https://www.graphpad.com |
| MOLREP (CCP4 suite) | (Potterton *et al.*, 2018) | www.ccp4.ac.uk |
| CCP4i2 Suite | (Potterton *et al.*, 2018) | www.ccp4.ac.uk |
| Chimera | (Pettersen *et al.*, 2004) | https://www.cgl.ucsf.edu/chimera/ |
| Coot | (Emsley *et al.*, 2010) | https://www2.mrc-lmb.cam.ac.uk/personal/pemsley/coot/ |
| CCP4 MG | (McNicholas *et al.*, 2011) | www.ccp4.ac.uk |
| Protein Thermal Shift™ Software v1.3 |  | Applied Biosystems™ |
| AKTA Unicorn 7 Evaluation software |  | https://www.cytivalifesciences.com/ |
| CLC viewer |  | http://resources.qiagenbioinformatics.com/manuals/clcsequenceviewer/current/index.php?manual=Introduction_CLC_Sequence_Viewer.html |
| JalView2 | (Waterhouse *et al.*, 2009) | www.jalview.org |
| Software used to fit urea denaturation data | (Perez-Riba and Itzhaki, 2017) |  |
| Chiascan pro-data viewer | Applied Photophysics |  |
| **Other** |  |  |
| CLARIOStar Plate Reader | BMG Labtech |  |
| CLARIOStar tryptopan excitation filter (295nm) | BMG Labtech |  |
| CLARIOStar tryptophan emission filter (360 nm) | BMG Labtech |  |
| ÄKTA pure chromatography system | Cytiva |  |
| CD Spectrophotometer Chirascan | Applied Photophysics |  |
| PHERAstar FS | BMG Labtech |  |
| PHERAstar FS excitation filter (337 nm) | BMG Lab tech |  |
| PHERAstar FS emission filter (620/665 nm) | BMG Labtech |  |

### Resource availability

### Lead Contact

Further information and requests for resources and reagents should be directed to and will be fulfilled by the Lead Contact, Laura Itzhaki.

### Materials availability

Plasmids for the expression of nanobodies NB06 and NB09, and p16 wild-type and mutant proteins are available from the lead contact.

### Data and code availability

All crystallographic results were deposited to and are accessible from the Protein Data Bank (PDB) https://www.rcsb.org/. p16INK4a-NB09 (PDB: 7OZT).

PDB accession codes for structures used within, but not derived from, this study: CDK6/p16INK4a (PDB: 1BI7) and GFP:GFP nanobody complex (PDB: 3OGO).

### Experimental model and subject details

Nanobody production: Nanobodies used in this study were prepared following the protocols described in (Pardon et al., 2014).

*In vitro* studies: Recombinant proteins used in this study were either expressed from recombinant *E. coli* (Lemo21(DE3)), TG1, Rosetta2 (DE3) pLYS-S, BL21 Star™ (DE3) or baculoviral infected *Spodoptera frugiperda* cells (Sf9) encoded on vectors listed in the Key Resources Table.

Cell line studies: Cell-based assays were carried out in Human embryonic kidney (HEK) 293T cells.

## Methods

### High-throughput p16 construct generation and expression screening

A gene encoding *E. coli* codon optimised full-length wild-type p16 flanked by a 5’ *BamHI* site and a 3’ *HindIII* site was designed, commercially synthesised and delivered in a pMA vector. High-throughput generation and expression screening of p16 constructs was performed according to methods described in Berrow *et al*, (Berrow et al., 2007).

PCR primers were used to amplify the wild-type p16 sequence. Each of the primers included 5’ extensions to enable high-throughput InFusion cloning into five different vectors which allowed the simultaneous addition of five different solubility tags: an N-terminal (pOPINF) or C-terminal (pOPINE) hexa-histidine-tag, or in addition to an N-terminal hexahistidine tag an N-terminal glutathione S-transferase (pOPINJ), maltose binding protein (pOPINM), or small ubiquitin modifying protein (pOPINS) tag. Plasmids were screened by transforming into chemically competent OmniMaxII cells before plating onto LB agar supplemented with 50 μg/mL carbenicillin, 100 μg/mL 5-bromo-4-chloro-3-indoyl-beta-D-galactopyranoside (X-Gal), 1 mM isopropyl beta-D-1-thiogalactopyranoside (IPTG). Selected colonies were cultured, plasmids were extracted and sequenced using a T7F sequencing primer to confirm the presence of the correct sequence. Constructs were then screened in *E. coli* Lemo21 (DE3) and in Rosetta2 (DE3) pLYS-S, purified by robotic immobilized-metal affinity chromatography (IMAC) and soluble p16 expression analyzed by SDS-PAGE. High levels of soluble expression were observed in *E. coli* Lemo21 (DE3) cells from the pOPINF plasmid. As such N-terminal histidine tagged p16 was chosen to take forward for further work. To generate p16^L37S/L121R^ inverse site-directed mutagenesis (SDM) was performed in two rounds of mutagenesis using the wild-type sequence as a template. Pairs of phosphorylated inverse primers with the codons encoding the mutation being present at the 5’ end of either of the primers (Table S4) were used to amplify the entire plasmid.

The resulting PCR product was treated with DpnI overnight at 37 °C to remove template DNA. Amplified product was purified, ligated and then transformed into chemically competent DH5α cells. The construct was verified by sequencing.

### Expression and purification of wild-type p16 and stabilising mutants

p16 wild-type and stabilising mutants were expressed by transforming plasmids into Lemo21(DE3) cells. *E. coli* was cultured in 2x TY media supplemented with 50 μg/mL carbenicillin and 35 μg/mL chloramphenicol in shake flasks at 37 °C and 225 rpm.

Once an optical density at 600 nm (OD_600_) of 0.6 was reached, cultures were thermally equilibrated to 20 °C and protein expression induced with the addition of 0.5 mM IPTG. Cultures were subsequently incubated overnight at 20 °C before harvesting by centrifugation at 12500 x g for 10 min. Cell pellets were resuspended in 35 mL lysis buffer (50 mM Tris-HCl pH 8.0, 500 mM NaCl, 20 mM imidazole, 1 mM dithiothreitol (DTT) supplemented with EDTA-free protease inhibitor tablets. Samples were subsequently homogenised and lysed by passing three times through an Emulsiflex-C5 cell disruptor at 100 mPa. The resulting lysate was centrifuged twice at 40000 x g for 30 min at 4 °C. Clarified lysates were loaded onto a 5 mL HisTrap FF Excel column at 2 mL/min using an AKTA pure chromatography system. The column was washed with 20 column volumes (CV) of wash buffer (50 mM Tris-HCl pH 8.0, 500 mM NaCl, 20 mM imidazole, 1 mM DTT) prior to protein being eluted by the addition of five CV of elution buffer (50 mM Tris-HCl pH 8.0, 500 mM NaCl, 500 mM imidazole, 1 mM DTT). Protein was immediately filtered through a 0.22 μm filter prior to loading directly onto a 16-600 G75 size-exclusion column pre-equilibrated with 20 mM Tris-HCl, pH 8.0, 200 mM NaCl, 1 mM DTT) at a flow rate of 1 mL/min. Protein was eluted at a constant flow rate. The purity of the eluted protein was evaluated by Coomassie-stained SDS-PAGE. Fractions corresponding to pure p16 (wild-type or p16^L37S/L121R^) were pooled and concentrated on VivaSpin concentrators with a 5 kDa molecular weight cut off.

### Expression and purification of cancer-associated mutant p16 proteins

For each p16 mutant, 100 ng of DNA (list in key resources table) was transformed into chemically competent Lemo21(DE3) *E. coli* cells, plated onto LB agar plates supplemented with 50 μg/mL carbenicillin and 35 μg/mL chloramphenicol and grown overnight at 37 °C. An overnight pre-culture was set up by inoculating a few colonies into 10 mL 2xTY media supplemented with 50 μg/mL carbenicillin and 35 μg/mL chloramphenicol and 1 % w/v glucose and growing overnight at 37 °C, 225 rpm.

Cultures were pelleted by centrifugation at 4000 x g for 10 min and then the supernatant discarded before resuspending in 1 L of 2xTY supplemented with 50 μg/mL carbenicillin and 35 μg/mL chloramphenicol. Cultures were grown at 37 °C, 225 rpm until an OD_600_ 0.6 was reached. Cultures were subsequently induced with 0.5 mM IPTG and grown for a further 16 hr at 20 °C shaking. *E. coli* cultures were harvested by centrifugation at 12500 x g for 10 min and pellets stored frozen at −20 °C. Pellets of *E. coli* expressing p16 mutants were defrosted and resuspended in 25 mL lysis buffer supplemented with EDTA-free protease inhibitor tablets (SigmaAldrich), 2 mg/mL lysozyme and 2 mg/mL DNase I. Cells were homogenised by passing three times through an Emulsiflex-C5 cell disruptor at 100 mPa. Cell debris and inclusion bodies were pelleted by centrifuging at 40,000 x g for 30 min before the supernatant was discarded. Inclusion bodies (IBs) were washed three times using the following method, initially each IB pellet was resuspended in 30 mL wash buffer 1 (50 mM Tris-HCl pH 8.0, 500 mM NaCl, 10% Triton-X100, 5 mM DTT) and sonicated for 3 x 15 sec on ice before centrifugation again at 40,000 x g for 30 min. IBs were washed in buffer 2 (50 mM Tris-HCl pH 8.0, 2 M NaCl, 1% Triton X-100, 5 mM DTT) to remove bound DNA, sonicated and pelleted. The IBs were subsequently washed in buffer 3 (50 mM Tris-HCl pH 8.0, 500 mM NaCl, 1 mM DTT) to remove detergent, sonicated and pelleted again. Lastly, IBs were resuspended in urea binding buffer (50 mM Tris-HCl pH 8.0, 500 mM NaCl, 20 mM imidazole, 8 M urea, 1 mM DTT) and left to solubilise at room temperature for 1 hr before insoluble material was removed by centrifugation for 30 min at 40,000 x g. Solubilised inclusion bodies were incubated with 1 mL of Ni-NTA beads for 1 hr at room temperature to allow binding to occur. Beads were pelleted, supernatant removed, and beads subsequently washed in 15 mL urea binding buffer. This wash step was repeated a further two times to remove non-specifically bound protein. p16 mutants were eluted from beads by the addition of 5 mL urea elution buffer (50 mM Tris-HCl pH 8.0, 500 mM NaCl, 500 mM imidazole, 8 M urea, 1 mM DTT). Eluted protein was quantified by nanodrop and flash frozen in 2 mL aliquots for long term storage at −20 °C.

### Expression and purification of Avi-tag-p16 protein

Human full length p16 (residues 1-166) was cloned into pGEX-6P-1 (Cytiva) in *BamH*I and *EcoR*I restriction sites to a generate a N-terminal glutathione-S-transferase (GST) tag followed by a 3C protease recognition site using a synthesised and codon optimised DNA template (Eurofins genomics). To generate GST-3C-Avi-tag p16, a N-terminal Avi tag was introduced using the QuikChangeII site-directed mutagenesis kit (Agilent). All constructs were verified by sequencing. p16 proteins were expressed in recombinant BL21STAR™ (DE3) *E. coli* chemically competent cells (ThermoFisher scientific) grown at 30 °C in 2xYT medium till OD_600_ ∼0.7-0.8, cooled, and induced at 18 °C by addition of 0.5 mM IPTG and then incubated for 16 hr. Cells were harvested by centrifugation (4000xg for 20 min at 4 °C), resuspended in buffer containing 10 mM HEPES pH 7.5, 150 mM NaCl and 0.5 mM TCEP supplemented with cOmplete EDTA-free protease inhibitor cocktail (Roche), 2 mg/mL DNase I and 5 mM MgCl_2_ and frozen prior to purification. Once thawed on ice, cells were lysed using sonication for total of 5 min with 30 % amplitude and pulsed for 20 s on and 40 s off. The lysate was subsequently cleared by centrifugation for 1 hr at 100,000 x g at 4 °C. Supernatant was recovered and incubated at 4 °C with shaking for 3 h with Glutathione Sepharose 4B resin (Cytiva), pre-equilibrated in 10 mM HEPES pH 7.5; 150 mM NaCl and 0.5 mM TCEP. The Glutathione Sepharose 4B resin was then loaded into an empty gravity flow column, the unbound fraction collected and the resin washed with 50 mL of buffer 10 mM HEPES pH 7.5; 150 mM NaCl and 0.5 mM TCEP. Bound protein was eluted in 1 ml fractions with buffer supplemented with 20 mM reduced glutathione at pH 8.0. Fractions containing protein were combined and incubated overnight using 1:50 (w/w) 3C protease at 4 °C. Cleaved protein was further purified by size exclusion chromatography (Superdex75 HiLoad (Cytiva) equilibrated in 10 mM HEPES pH 7.5; 150 mM NaCl and 0.5 mM TCEP. Selected fractions were analysed by SDS-PAGE, pooled and flowed through Glutathione Sepharose 4B resin. Unbound protein was concentrated to 5 mg/ml, flash frozen using liquid nitrogen and stored at −80 °C.

### Avi-tagged p16 biotinylation *in vitro*

Avi-p16 (40 μM) was incubated with 10 μg BirA in biotinylation buffer (50 mM Bicine pH 8.3, 10 mM ATP, 10 mM MgOAc, 50 μM *d*-biotin) at 30 °C for 30 min. The protein mixture was then buffer exchanged using size exclusion chromatography into HTRF buffer A (50 mM HEPES pH 7.5, 100 mM NaCl). The extent of protein biotinylation was monitored by mass spectrometry.

### Expression and purification of CDK4 and CDK6

Human CDK4 and CDK6 full length (residues 1-303 and residues 1-326, respectively) were expressed in *Sf*9 insect cells using a recombinant baculovirus expression system (Bieniossek et al., 2012), Geneva-Biotech MultiBac™. Initially to generate an N-terminal glutathione-S-transferase (GST-tag) followed by a 3C protease recognition site full length human CDK4 or CDK6 were cloned into a pGEX6P1vector (Cytiva) at the *BamH*I and *Xho*I restriction sites. GST-CDK4 and GST-CDK6 constructs were subsequently subcloned into the pACE-BAC1 acceptor vector at the *BamH*I and *EcoR*I restriction sites. All constructs were verified by restriction digestion and DNA sequencing.

Approximately ∼ 1 µg of an acceptor vector, pACE-Bac1-GST-CDK4 or pACE-Bac1-GST-CDK6, were used to transform DH10*EmBacY E. coli* cells harbouring the *EmBacY* MultiBac™ bacmid and subsequently screened by blue/white selection on plates containing kanamycin, gentamycin, and tetracycline. White recombinant colonies were selected for bacmid preparation. One white colony was grown overnight in 2 mL LB media at 37 °C shaking at 160 rpm and MultiBac bacmid DNA was prepared by alkaline lysis. The bacterial cells were centrifuged at 4000 x g for 10 min and the media removed. Pelleted cells were lysed and cell precipitates removed using the Qiagen Miniprep kit buffers P1, P2 and N3. The bacterial pellet was resuspended in 300 µL of P1 buffer, 300 µL of Buffer P2 was added and mixed gently by inverting the tube to generate a homogenous mixture. 300 µL of N3 buffer was then added and the mixing step repeated prior to centrifugation for 10 min at 16,000 x g. The supernatant containing bacmid DNA was recovered into a new Eppendorf, precipitated using 40 % isopropanol and subsequently washed twice using 70 % ethanol prior to resuspension in 50 µL of sterile water and 300 µL Insect-Xpress insect cells medium (Lonza) in a sterile hood. 10 µL GeneJuice® Transfection reagent (Merck Millipore) was then added to the DNA before adding it to the *Sf*9 cells (3 mL at (0.7-1)×10^6^ cells per ml) in a 6 well plate for 72 hr. Infection efficiency was assessed by the presence of YFP-expressing cells. Harvested virus V0, present in the media of the infected cells, was subsequently added to 50 mL of fresh *Sf*9 cells at (0.7-1) x10^6^ cells per mL and grown for 65-72 hr before harvesting the next generation of virus (V1). This step was repeated to obtain virus V2, which was used for recombinant protein expression. For protein expression (0.7-1) x10^6^ cells per mL were infected with the V2 viral stock for 65-72 hr. The amount of added virus was optimised to allow one round of cell duplication, after which cell division stopped and cell metabolism was switched to allow protein expression without further cell duplication. Cells were harvested by centrifugation (4000 x g for 20 min at 4°C), resuspended in buffer containing 10 mM HEPES pH 7.5, 150 mM NaCl and 0.5 mM TCEP supplemented with EDTA-free protease inhibitor cocktail (Roche), 2 mg/mL DNase I and 5 mM MgCl_2_ and frozen prior to purification. To purify proteins, pellets of insect cells (following GST-CDK4 or GST-CDK6 expression) were thawed on ice and then lysed by sonication for total of 5 min with 30% amplitude and pulsed for 20 s on and 40 s off. Lysate was subsequently cleared by centrifugation for 1 hr at 100,000 x g at 4 °C. The supernatant was then incubated for 3 h at 4 °C with shaking with glutathione Sepharose 4B resin (Cytiva), pre-equilibrated in 10 mM HEPES pH 7.5, 150 mM NaCl and 0.5 mM TCEP. The glutathione Sepharose 4B resin was then loaded into an empty gravity flow column, the unbound fraction was collected and then the resin washed with 50 mL of buffer (10 mM HEPES pH 7.5, 150 mM NaCl and 0.5 mM TCEP). Bound proteins were eluted in 1 ml fractions with buffer supplemented with 20 mM reduced glutathione at pH 8.0. GST-CDK6 (or GST-CDK4) containing fractions were combined, incubated overnight with 1:50 (w/w) 3C protease at 4 °C. Cleaved and uncleaved CDK6 (or CDK4) were further purified by size exclusion chromatography (Superdex200 HiLoad (Cytiva)) equilibrated in 10 mM HEPES pH 7.5, 150 mM NaCl and 0.5 mM TCEP. Fractions containing cleaved protein were re-applied to glutathione Sepharose 4B resin to remove the GST and 3C prior to concentrating. Fractions containing uncleaved proteins were collected and concentrated. All proteins (cleaved and uncleaved) were concentrated to *circa* 5 mg/ml, flash frozen using liquid nitrogen and stored at −80 °C.

### Immunisations and library construction

Immunisations and library construction were performed by Dr Els Pardon and co-workers as described (Pardon et al., 2014). Two libraries were generated by immunising *llama glama* with either p16 wild-type (Library 173) or the stabilized double mutant p16^L37S/L121R^ (Library 174). The resulting library of nanobodies for each library was cloned into a pMESy4 phage display vector which resulted in the nanobody gene fused to a C-terminal histidine tag, an EPEA capture select detection tag followed by an amber stop codon, a Hemagglutinin (HA) tag and finally the pIII phage coat protein. The entire library was transformed into TG1 *E. coli* cells and grown to an OD_600_ of 0.6 in 50 mL 2xTY media supplemented with 100 μg/mL ampicillin and 1 % w/v glucose. The culture was pelleted by centrifugation, then resuspended in 5 mL 2xTY supplemented with 20 % v/v glycerol and frozen as a glycerol stock at −80 °C until required.

### Nanobody phage library rescue

In two separate baffled 250 mL flask, 60 mL of 2xTY supplemented with 10 μg/mL ampicillin and 2 % w/v glucose was inoculated respectively with six OD_600_ units of glycerol stock for each phage display. Each flask was grown at 37 °C, 200 rpm until an OD_600_ 0.5 was reached whereby 10 mL of each culture was transferred to a new 50 mL falcon tube and infected with 4.0 x 10^10^ plaque forming units (pfu) of VCSM13 helper phage. Tubes were incubated at 37 °C in a static incubator prior to bacteria being pelleted by centrifuging for 10 minutes at 2800 x g. The supernatant was removed and discarded before resuspending cells. These were used to inoculate 50 mL 2xTY media supplemented with 100 μg/mL ampicillin, 25 μg/mL kanamycin and incubated at 37 °C, 200 rpm overnight.

### Phage precipitation

To precipitate phage, cells were centrifuged at 3200 x g for 10 min at 4 °C. To a fresh falcon 40 mL of supernatant containing phage particles was transferred and mixed by inversion with 10 mL of 20 % w/v poly(ethylene glycol) molecular weight 6000 (PEG6000), 2.5 M NaCl before incubating on ice for 30 min. Precipitated phage were pelleted by centrifuging at 4 °C for 10 min at 2300 x g prior to the supernatant being removed and discarded. The precipitated phage pellet was resuspended in 1 mL of ice-cold phosphate buffered saline and centrifuged at 20,000 x g, 4 °C for 1 min. To a fresh tube, 1 mL of the supernatant was transferred and precipitated again, by the addition of 250 μL 20 % w/v PEG6000, 2.5 M NaCl, mixed by inversion and incubated on ice for a further 10 min. Precipitated phage were centrifuged at 20,000 x g for 15 min, supernatant removed and pelleted phage resuspended in 1 mL ice-cold PBS. Phage were centrifuged again for 15 min and finally the supernatant transferred to a fresh tube.

### Phage library titration

To determine the number of infective phage in each library, a library titration was performed. Serial dilutions of each library were performed by mixing 10 μL of phage with 90 μL of PBS. A 10 μL sample of each dilution was transferred onto 90 μL of mid-log phase TG1 *E. coli* cells (OD_600_ 0.5) and incubated in a static incubator at 37 °C for 15 min to infect cells. After infection 5 μL of infected cells were transferred onto LB agar, 100 μg/mL ampicillin, 2 % glucose plates, left to air dry for 20 min and then grown overnight at 37 °C. The number of infective phage was determined using the following formula:

Infective phage = (Number of colonies x 200 x 10 x dilution factor)

### Nanobody phage display rounds 1 and 2

Six different panning strategies were explored for each library to maximise the chance of obtaining unique binders to p16: (i) solid phase coated wild-type p16; (ii) solid phase coated p16^L37S/L121R^; (iii) solid phase coated wild-type p16 and CDK6 complex; (iv) solid phase coated stabilized p16 and CDK6 complex; (v) CDK6 capture of p16 wild-type; and (vi) neutravadin capture of biotinylated p16. Briefly, 50 μg/mL of protein for round 1 and 5 μg/mL for round 2 was diluted in PBS for each panning strategy. For each, 100 μL was added to a well on a Nunc Maxisorp high binding 96-well plate along with a control. Plates were incubated overnight at 4 °C to allow passive adsorption to occur. Wells were emptied and washed five times with 250 μL PBS prior to the addition of 4 % w/v non-fat dried milk in PBS (MPBS) to block wells. Wells were blocked for two hr at room temperature prior to washing again. In the case of the two capture panning strategies a further step involving the addition of wild-type p16 or biotinylated p16 was added. Hereby, 50 μg/mL of the respective protein was diluted in 4% MPBS and added to the plates for a further 2 hr after blocking to allow binding. Wells were washed again before proceeding. To each well 90 μL of 1 % MPBS was added, followed by 10 μL of output phage. Phage were allowed to bind for 2 hr shaking at 700 rpm prior to extensive washing. Wells were washed five times with 250 μL PBS supplemented with 0.2 % v/v tween-20 (PBST) followed by a further ten times with PBS only. Phage particles were eluted by the addition of 100 μL solution of 0.25 mg/mL trypsin in PBS and incubated for 30 min to elute protein. Trypsin activity was inhibited by the addition of 5 μL of 4-(2-aminoethyl)benzenesulfonyl fluoride hydrochloride (AEBSF) at 4 mg/mL.

Phage titrations were performed as previously described, by serial titrating 10 μL of phage into 90 μL of PBS and used to infect 90 μL of exponential growing TG1 cells.

### Phage repertoire rescue

For each library, 50 μL of output phage was used to infect 1 mL of exponentially growing TG1 *E. coli* cells. The phage were allowed to infect for 30 minutes at 37 °C with no shaking. Subsequently 5 mL of 2xTY supplemented with 100 μg/mL ampicillin, 2 % w/v glucose was added and grown overnight at 37 °C, 200 rpm. A glycerol stock of this was prepared by mixing equal amounts of culture with 50% glycerol in 2xTY and snap freezing at −80 °C for long term storage. To rescue phage repertoire for subsequent rounds of phage display a 1/1000 dilution of overnight culture was made in 25 mL 2xTY supplement with 100 μg/mL ampicillin and 2 % w/v glucose and grown at 37 °C, 200 rpm until an OD_600_ 0.5 was reached. To 10 mL of this culture, 2×10^10^ pfu of VSCM13 helper phage were added and culture mixed by gentle inversion. Helper phage were allowed to infect for 30 min at 37 °C with no shaking. Cells were pelleted by centrifuging at 2800 x g for 10 min and supernatant removed and discarded. The cell pellet was resuspended in 50 mL 2xTY supplemented with 100 μg/mL ampicillin, 25 μg/mL kanamycin and grown overnight at 37 °C, 200 rpm to allow phage expression. Phage particles were rescued as previously described.

### Monoclonal nanobody expression

Forty microlitres of 10-fold serial dilutions of TG1 *E. coli* cells were plated on LB selection plates (LB agar, 100 μg/mL ampicillin, 2 % w/v glucose) and grown overnight at 37 °C inverted. Single colonies were used to inoculate a master plate with each well containing 100 μL 2xTY supplemented with 100 μg/mL ampicillin, 2 % w/v glucose, 10% w/v glycerol and grown overnight at 37 °C shaking. The plate was frozen at −80 °C for long-term storage. To each well of a 96-well deep well block 1 mL of 2xTY supplemented with 100 μg/mL ampicillin, 0.1 % w/v glucose was added. Using a multichannel pipette 10 μL of overnight master plate culture was added, blocks covered with gas permeable adhesive seal and grown at 37 °C, 200 rpm until an OD_600_ 0.6 was reached. To induce nanobody expression 1 mM IPTG was added to each well. Cells were incubated for a further four hr at 37 °C prior to being pelleted by centrifuging block at 3200 x g for 10 min. Supernatant was discarded and cell pellets containing periplasmically expressed nanobodies were frozen at −20 °C until required.

### Periplasmic extraction

Expression culture blocks were thawed at room temperature prior to the addition of 100 μL PBS to each well. Blocks were shaken at 1200 rpm for 30 min to release nanobodies from the periplasm. To pellet cells, blocks were spun for 10 min at 3200 x g, 4 °C before 90 μL of periplasmic fraction was transferred to a fresh 96-well plate.

### Monoclonal phage Enzyme Linked Immunosorbant Assay (ELISA) and DNA isolation

An ELISA was performed by coating a Nunc Maxisorp plate with 5 μg/mL antigen (in the same format that they were coated for biopanning). The plate was washed five times with 200 μL PBST between all subsequent steps. The plate was subsequently blocked by incubating with 200 μL 4% w/v MPBS for 1 hr before washing. To each well 20 μL of periplasmic fraction was diluted in 80 μL 1% w/v MPBS and incubated for 2 hr. Wells were washed and then 100 μL of secondary antibody solution (containing a 1:1000 dilution of streptavidin alkaline phosphatase (Promega) and a 1:4000 dilution of Capture Selection C-tag (ThermoFisher) diluted in 4% w/v MPBS) was added to each well.

Plates were incubated for 1 hr prior to washing as previous. A 2 mg/mL solution of ρ-nitrophenyl phosphate (pNPP) was then added and incubated at room temperature before reading at 405 nm. Of the positive binders, 192 were randomly selected and reformatted into a new masterplate by inoculating 1 mL of Agencourt Broth (Beckman Coulter) supplemented with 100 μg/mL ampicillin and 2% w/v glucose with glycerol stock for each and grown overnight at 37 °C. A glycerol stock of this plate was produced before the masterplate cultures were pelleted by centrifuging at 3200 x g for 10 min, supernatant disposed of and frozen at −20 °C. Using a CosMCPrep robotic platform (Beckman Coulter) a miniprep was performed on all samples. DNA samples were sent for sequencing and the sequences were loaded into CLC viewer, aligned, and trimmed to remove extra base reads. The sequences were translated and the CDR3 sequence extracted. A pairwise comparison was performed on this region and sequences subsequently aligned according to these families.

### Large-scale nanobody expression

Nanobodies were expressed in the periplasm of Rosetta2 *E. coli* cells from the pMESy4 vector that appends a C-terminal histidine tag followed by the four amino acid (EPEA) CaptureSelect C-tag sequence (ThermoFisher). Briefly, chemically competent Rosetta2 *E. coli* were transformed by heat-shock transformation using 1 μL of nanobody pMESy4 plasmid. Transformed cells were plated onto LB agar supplemented with 100 μg/mL carbenicillin, 35 μg/mL chloramphenicol and 1% w/v glucose and grown overnight at 37°C. A 10 mL overnight starter culture was set up for each construct using 5 mL 2xTY media supplemented with 100 μg/mL carbenicillin, 35 μg/mL chloramphenicol and 1% w/v glucose and grown at 37 °C overnight, 225 rpm. Overnight cultures were pelleted by centrifuging at 4000 x g for 5 min, supernatant discarded and pelleted cells resuspended in 1 L of 2xTY media supplemented with 100 μg/mL carbenicillin, 35 μg/mL chloramphenicol, 0.1 % w/v glucose and 1 mM MgSO_4_. Large-scale cultures were grown at 37 °C until an OD_600_ 0.6 was reached. Cultures were thermally equilibrated to 28 °C and protein expression induced by the addition of 1 mM IPTG. Cultures were subsequently incubated overnight at 28 °C, 225 rpm before being harvested by centrifuging at 12500 x g for 10 min. Pellets were frozen at −20 °C prior to purification.

### Periplasmic release of nanobodies using cold-osmotic shock

The periplasmic fraction containing the expressed nanobodies was released by spheroplasting cells using cold osmotic shock. Cell pellets were defrosted and resuspended overnight in 15 mL of TES buffer (0.2 mM Tris-HCl pH 8.0, 0.5 M sucrose, 1 mM EDTA supplemented with an EDTA-free protease inhibitor tablet (Sigma)) by shaking at 4 °C, 225 rpm. Periplasmic release was achieved by the addition of 30 mL of ice-cold TES4 buffer (40 mM Tris-HCl pH 8.0, 0.1 M sucrose, 200 μM EDTA supplemented with 2 mg/mL DNase I) before shaking for a further hour at 4 °C. Cell debris was removed by centrifuging for 10 min at 12500 x g before further clarification of the periplasmic fraction by centrifuging at 40000 x g for 30 min at 4 °C.

### Nanobody purification

Cleared periplasmic fractions were loaded onto a 1 mL HisTrap FF Excel column at 2 mL/min using an AKTA pure chromatography system (both Cytiva Life Sciences). Upon complete loading the column was washed with 20 CV of wash buffer (50 mM Tris-HCl pH 8.0, 500 mM NaCl, 20 mM imidazole) at a flow rate of 2 mL/min. Protein was subsequently eluted by the addition of 5 CV of elution buffer (50 mM Tris-HCl pH 8.0, 500 mM NaCl, 500 mM imidazole) in a reversed flow direction at 2 mL/min.

Eluted protein was filtered through a 0.22 μm filter prior to loading directly onto a 16-600 G75 gel-filtration column pre-equilibrated with size exclusion buffer (50 mM Tris-HCl pH 7.5, 100 mM NaCl) at a flow rate of 1 mL/min. Protein was eluted at a constant flow rate. The peak corresponding to the nanobody was collected as 2 mL fractions. Eluted protein was analysed by running a sample of each nanobody on a 15 % SDS-PAGE gel and staining with Coomassie G250 stain to confirm purity. Nanobody fractions were pooled and concentrated using a 5000 MWCO VivaSpin centrifuge concentrator.

Concentrated protein was aliquoted and snap frozen in liquid nitrogen prior to long term storage at −80 °C.

### Circular dichroism (CD) and thermal denaturation of nanobodies

Each nanobody construct was dialysed into 20 mM Tris-HCl pH 7.5, 50 mM NaCl overnight at 4 °C, then diluted to a concentration of 10 μM and incubated at 20 °C for 1 hr. Using a 1 mm path length quartz cuvette, 300 μL of protein was added and allowed to equilibrate to 20 °C for 5 minutes prior to signal acquisition. Far-UV CD data were collected from 200-280 nm in triplicate. The data were averaged and then converted to mean residue molar ellipticity. The temperature of the sample was subsequently increased to 95 °C at a ramp rate of 1 °C/min, and the ellipticity recorded at 218 nm.

Upon reaching the final temperature a CD spectrum was again acquired. Subsequently, the temperature of each sample was cooled to 20 °C at a rate of 1 °C/min. Upon reaching 20 °C a final CD spectrum was acquired to test for the reversibility of unfolding.

### Urea denaturation of nanobodies

For the following experiments urea concentration was determined at 25 °C by measuring the change in refractive index between urea buffer and non-urea buffer. The protein concentration of each nanobody was determined using the nanodrop prior to being diluted to a final concentration of 20 μM in 50 mM Tris-HCl pH 7.5, 100 mM NaCl. Using a Hamilton microtitrator dispenser a 46-point linear titration of urea was dispensed into a Corning 96 well low-binding solid black assay plate by dispensing appropriate volumes of non-urea buffer (50 mM Tris-HCl pH 7.5, 100 mM NaCl) and urea buffer (50 mM Tris-HCl pH 7.5, 100 mM NaCl, 9 M urea) in a final volume of 180 μL. A replicate plate was also prepared, using the same buffers but with the addition of 1 mM DTT. To each well 20 μL of each nanobody was dispensed, plates sealed with an aluminium plate seal and shaken for 5 min on an orbital shaker at 800 rpm to mix completely. Plates were incubated at 25 °C for 1 hr before tryptophan fluorescence was measured by reading on a CLARIOstar plate reader. Excitation wavelength was at 295 ± 10 nm and emission wavelength at 360 ± 20 nm. Data were fitted to a two-state model.

### Urea denaturation of wild-type and mutant p16 in the presence of p16 nanobodies

Purified p16 wild-type, p16 mutants and nanobodies (NB03, NB05, NB06, NB09, NB16 and NB17) were buffer exchanged into 4 M urea buffer (50 mM Tris-HCl pH 7.5, 100 mM NaCl, 4 M urea, 1 mM DTT). For each protein, a centripure p25 buffer-exchange column (EMP Biotech) was pre-equilibrated with 25 mL of 4 M urea buffer before 2.5 mL of protein was added to the column and allowed to flow through by gravity.

Subsequently 3.5 mL of 4 M urea buffer was applied to each column and the eluted protein was collected. Protein concentration was determined, and then the samples were diluted to a final concentration of 40 μM. Equimolar amounts of p16 and each nanobody were mixed to a final volume of 4 mL. Using a Hamilton microtitrator dispenser a 48-point titration of urea with 0.1 M increments was dispensed into a Corning 96-well low-binding solid black assay plate by dispensing appropriate volumes of non-urea buffer (50 mM Tris-HCl pH 7.5) and urea-containing buffer (50 mM Tris-HCl pH 7.5, 100 mM NaCl, 5 M Urea, 1 mM DTT) in a final volume of 180 μL. To each well 20 μL of protein complex was dispensed on top, plates sealed with an aluminium plate seal and shaken for 5 min on an orbital shaker at 800 rpm to mix completely. Three curves were dispensed for each sample. Plates were incubated at 25 °C for 2 h before tryptophan fluorescence was measured by reading on a CLARIOStar plate reader. The excitation wavelength was 295 ± 10 nm and emission wavelength was 360 ± 20 nm.

Denaturation curves of the p16 mutants in the presence of nanobodies (NB06 and NB09) were performed in an identical manner to wild type as described above and fitted to a two-state model.

### Crystallisation of p16-nanobody complexes

Wild-type p16 (with an N-terminal hexahistidine tag, generated from the pOPINF construct) and each nanobody was concentrated to 300 µM in a 5 kDa MWCO VivaSpin concentrator. Each nanobody was then mixed with p16 at a molar concentration of 1.2:1 respectively and incubated on ice for 1 h prior to loading onto a 16-600 G75 size-exclusion chromatography column (Cytiva) pre-equilibrated with 20 mM Tris-HCl pH 7.5, 100 mM NaCl. Protein was eluted at a flow rate of 1 mL/min and fractions corresponding to A_280nm_ peaks were analysed by SDS-PAGE. Peaks corresponding to stable nanobody-p16 complexes were pooled and concentrated in a 5000 kDa MWCO VivaSpin concentrator to a final concentration of 10 and 14 mg/mL respectively for p16-NB06 and p16-NB09. Crystal screens of p16-nanobody complexes were performed in 96-well sitting-drop vapor diffusion plates at 18 °C. Crystals appeared for the p16-NB09 complex after 2-7 days in several different conditions across a variety of different commercial screens such as Index (Hampton Research), Morpheus or Proplex (Molecular Dimensions). Crystals grown directly from Morpheus condition G4 from Tube 2-28 consisting of carboxylic acids, buffer system I and Precipitant Mix 4 (Molecular Dimensions) were mounted in loops and cooled in liquid nitrogen before data collection at 100 K on Beamline I03, Diamond Light Source (DLS), Harwell, UK.

### Data collection and structure refinement

Data were collected at the Diamond Light Source (Oxfordshire, UK) on beamline I03 (MX13587). Data was processed using XIA2/DIALS (Winter et al., 2018) and POINTLESS/AIMLESS (Evans, 2011; Evans and Murshudov, 2013) run through the CCP4i2 GUI (Potterton et al., 2018; Winn et al., 2011). The p16-NB09 structure was solved by molecular replacement using Phaser (McCoy et al., 2007) and Buccaneer (Cowtan, 2012), and PDB entry 1BI7 (structure of CDK6-p16 complex) as the search model for p16 and 3OGO (structure of a GFP-GFP nanobody) as a search model for a nanobody. The structure was refined using cycles of automated refinement in REFMAC5 (Murshudov et al., 2011) and manual refinement in Coot (Emsley et al., 2010). The final structural parameters were evaluated using Molprobity (Williams et al., 2018) and Coot. All figures were produced using UCSF Chimera (Pettersen et al., 2004). The statistics of the data sets and the crystallographic refinement are presented in Table S3.

### Homogenous time-resolved fluorescence (HTRF)

A direct binding assay format was used to measure the affinity of p16 for CDK4 or CDK6 in the presence or absence of a nanobody. GST-tagged CDK4 or CDK6 was incubated with biotinylated Avi-tagged p16 or Avi-tagged p16-nanobody complex to form a GSTCDK-p16 (+/- nanobody) complex. The complex was then incubated with a Tb-labelled anti-GST antibody and streptavidin-tagged XL665 dye. Formation of a complex brings the Tb and XL665 into proximity so that excitation of the Tb results in emission from the XL665 dye as a result of Förster resonance energy transfer (FRET) between the two probes. For direct binding measurements the biotinylated protein of interest was initially titrated, over 11 serial dilution points and an additional buffer blank point, against 5 nM of either GSTCDK4 or GSTCDK6. Concentrations of GSTCDK and the binding protein of interest were prepared in HTRF buffer A (50 mM HEPES, 100 mM NaCl, 1 mM DTT and 0.1 mg/ml BSA) and incubated together for 60 min at 4 °C.

5 nM Tb labelled anti-GST antibody and SAXL665 at 1/8th the concentration of the biotinylated protein, were prepared in HTRF buffer B (50 mM HEPES, 100 mM NaCl and 0.1 mg/ml BSA) and added to each well. The plate was incubated for a further 60 min at 4 °C, before being scanned. Samples were excited using a wavelength of 337 nm and emission spectra measured at 620 nm and 665 nm (PHERAstar FS (BMG LABTECH)). Binding curves were plotted using GraphPad Prism 9 from which the K_d_s were determined. The curves shown are representative binding curves from at least two runs carried out on separate days.

### Surface plasmon resonance (SPR)

All SPR experiments were performed on a Biacore S200 (Cytiva) at 4 °C using SPR buffer (10 mM HEPES pH 7.5, 150 mM NaCl, 3 mM EDTA and 0.005% Tween 20). Samples were centrifuged at 10000 x g for 10 min at 4 °C before use. 100 μg/mL GSTp16 was captured on a surface of goat anti-GST antibody (Cytiva) at 10 μL/min for 240 s. Antibody capture to the CM5 BIAcore sensor chip was via amine coupling using the standard protocol provided in the GST capture kit (Cytiva). Analyte solutions of nanobody NB06 and nanobody NB09 at 500 nM, 167 nM, 55.6 nM, 18.5 nM, 6.20 nM and 0 nM were flowed over the bound GSTp16 at 30 μL/min for 240 s and the dissociation was measured over 600 s. CDK6 was used as a positive control and titrated as an analyte at 500 nM, 167 nM, 55.6 nM, 18.5 nM, 6.20 nM and 0 nM for 240 s and dissociation was measured at 600 s. The bound GSTp16 was removed from the antibody using 10 mM glycine pH 2.1 at 30 μL/min for 240 s. The chip was regenerated by capturing fresh GSTp16 before each analyte concentration run. GST was loaded onto a separate lane at 10 μg/mL for 180 s at 10 μL/min to act as a negative control.

Sensorgram readings from the GST control were subtracted from GSTCDK sensorgrams to correct for non-specific interactions and bulk effects. K_d_ values and dissociation rates were calculated using the Biacore S200 Evaluation Software (Cytiva).

### Isothermal titration calorimetry (ITC)

ITC was used to quantify the binding affinities of the nanobodies (NB06 and NB09) for p16. The release of heat upon binding was measured at 25 °C using a MicroCal VP instrument (Malvern). Prior to sample loading, the sample chamber and syringe were washed with water then buffer and the samples degassed (15 min). In total, 65 injection steps of 6 µL of nanobody (128–180 µM) were made into a p16 solution in the cell (18-19 µM). Data were fitted using the Origin software package (Microcal) and NITPIC (MBR) software

### Cell culture

Human embryonic kidney (HEK) 293T cells were cultured in Dulbeccos’s Modified Eagles’s Medium (Gibco) supplemented with 10 % foetal bovine serum (SigmaAldrich) and 1 % penicillin/streptomycin (SigmaAldrich) in a humidified incubator at 37 °C and 5% CO_2_.

### Nanobody expression in mammalian cells, co-immunoprecipitation and western blot

NB06 and NB09 were expressed in HEK293T cells from a modified pOPINF vector (Berrow et al., 2007). Briefly, pOPINF was linearised by digestion with FastDigest *NcoI* and *HindIII* (ThermoFisher) and then dephosphorylated using FastAP (Thermofisher) before gel purification. A DNA GeneBlock encoding the blue-white colony screening cassette with a downstream HA-tag flanked by an upstream *EcoRI* site, was ordered from GeneArt (Thermofisher). This DNA was double digested using FastDigest *NcoI* and *HindIII* enzymes (ThermoFisher), then heat inactivated before PCR purification and subsequent ligation back into the pOPINF vector. Modified vector DNA (pOPINF-HA) was confirmed by sequencing. To express nanobodies NB06 and NB09, pOPINF-HA was first linearised by digestion with FastDigest *NcoI* and *EcoRI*, subsequently the coding sequence for each nanobody was PCR-amplified with primers containing InFusion compatible overhangs on the 5’ end of each (sequences provided in Table S4). The PCR amplified construct was then cloned into the linearised vector.

Constructs (pOPINF-HA_NB09/NB06) were confirmed by sequencing. This cloning strategy resulted in the removal of the periplasmic targetting PelB leader sequence and concurrently replaced the constructs with a C-terminal HA-tag instead of the C-terminal Histidine tag-CaptureSelect tag (HHHHHH-EPEA).

HEK293T cells were seeded in 10 cm dishes (Corning) and grown to 50-80 % confluency before transfection (5 µg of pOPINF-HA_NB09/NB06) using Lipofectamine 2000 (3 µL/µg of DNA) (Thermo Fisher Scientific). 48 hr post-transfection, cells were washed with PBS, lysed in lysis buffer (25 mM Tris-HCl pH 7.5, 225 mM KCl and 1% NP-40) supplemented with protease inhibitors (Sigma-Aldrich) and 1mM Na_3_VO_4_, 10 mM NaF and 1 mM phenylmethylsulfonyl fluoride (PMSF). Cell lysates were centrifuged at 14,000 x g for 20 min at 4 °C and incubated with anti-HA beads (Sigma-Aldrich) on a rotating wheel for 4 hr at 4 °C. Beads were then washed five times with lysis buffer and protein complexes eluted using 2x Laemmli buffer. Samples were subsequently analysed by Western blot. Samples were resolved using SDS-PAGE and transferred onto a 0.45 µM Immobilon-P PDVF membrane (Millipore) according to standard protocols. Membranes were immunoblotted with antibodies against p16 (Abcam, Ab108349) and HA-tag (Cell Signalling Technology, 3724). After primary antibody incubation, membranes were probed with the polyclonal swine anti-rabbit immunoglobulins/HRP (Dako, P0399) secondary antibody, developed using an enhanced chemiluminescence detection reagent (GE Healthcare) and imaged using the Odyssey CLx Infrared Imaging System (Li-Cor).

## Quantification and statistical analysis

Details of the quantification and statistical analysis carried out for each technique are provided in the relevant Methods subsections and where appropriate in the accompanying Figure Legends.

## Abbreviations

CDK: cyclin-dependent kinase

