## Supplemental information for "Nanobodies restore stability to cancer-associated mutants of tumor suppressor protein p16^INK4a^"

**Short heading: Identification of nanobodies to stabilize p16$^{INK4a}$**

**Abbreviations:**

**CDK: cyclin-dependent kinase**

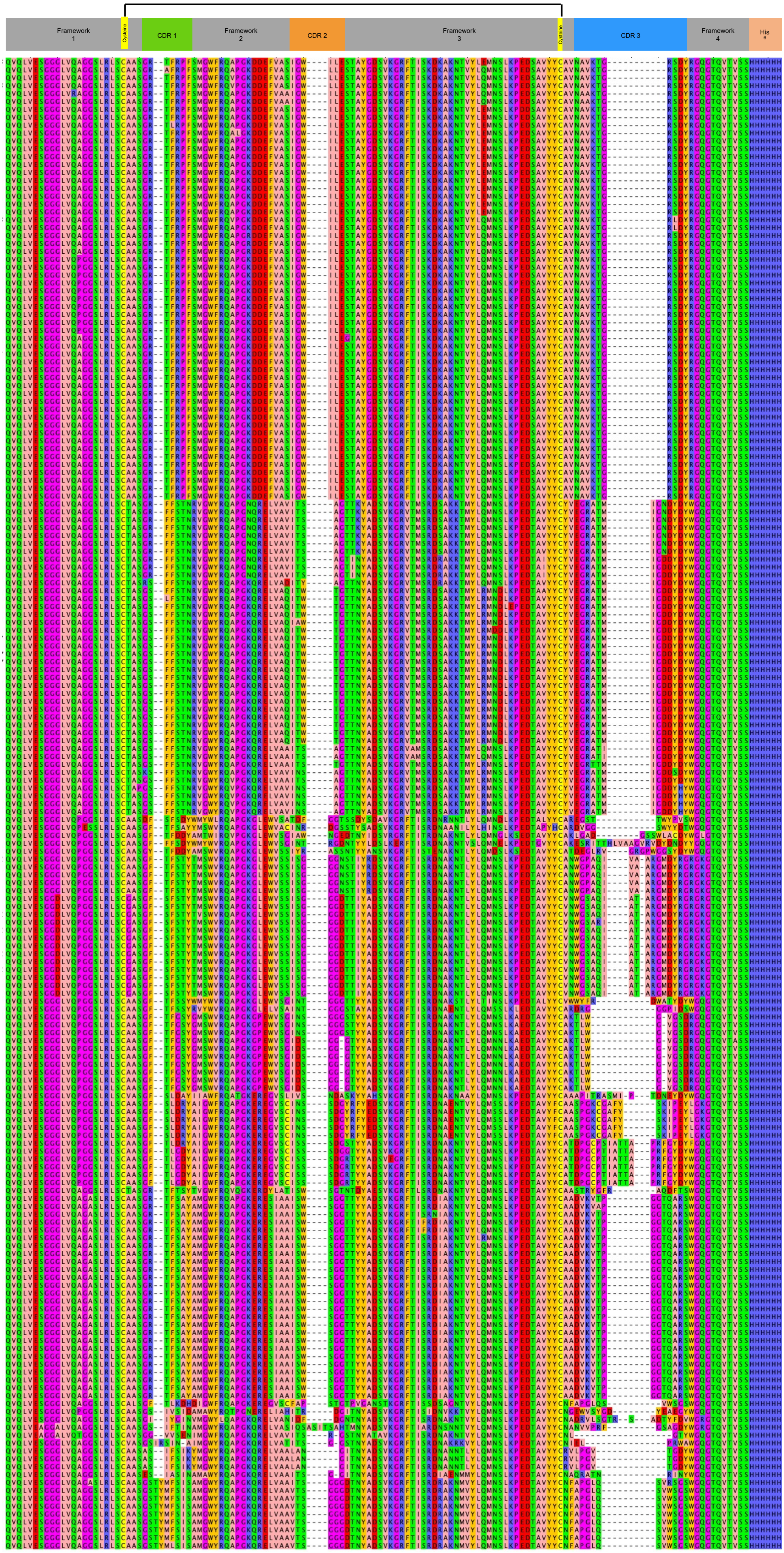
Framework 1
Cysteine
CDR 1
Framework 2
CDR 2
Framework 3
Cysteine
CDR 3
Framework 4
His 6

**Figure S1. Nanobody families and overall structure.** Related to Figure 1. Sequence alignment of all selected antibodies. Six large nanobody families dominated the phage display output: over 60% of the nanobodies belong to these families. Nanobodies within the same family are identified by CDR3 sequence alignment. Sixteen unique families consisted of only a few nanobody clones. The complementarity determining regions (CDR), CDR1, CDR2 and CDR3 are aligned. Figure generated using JalView (Waterhouse et al., 2009).

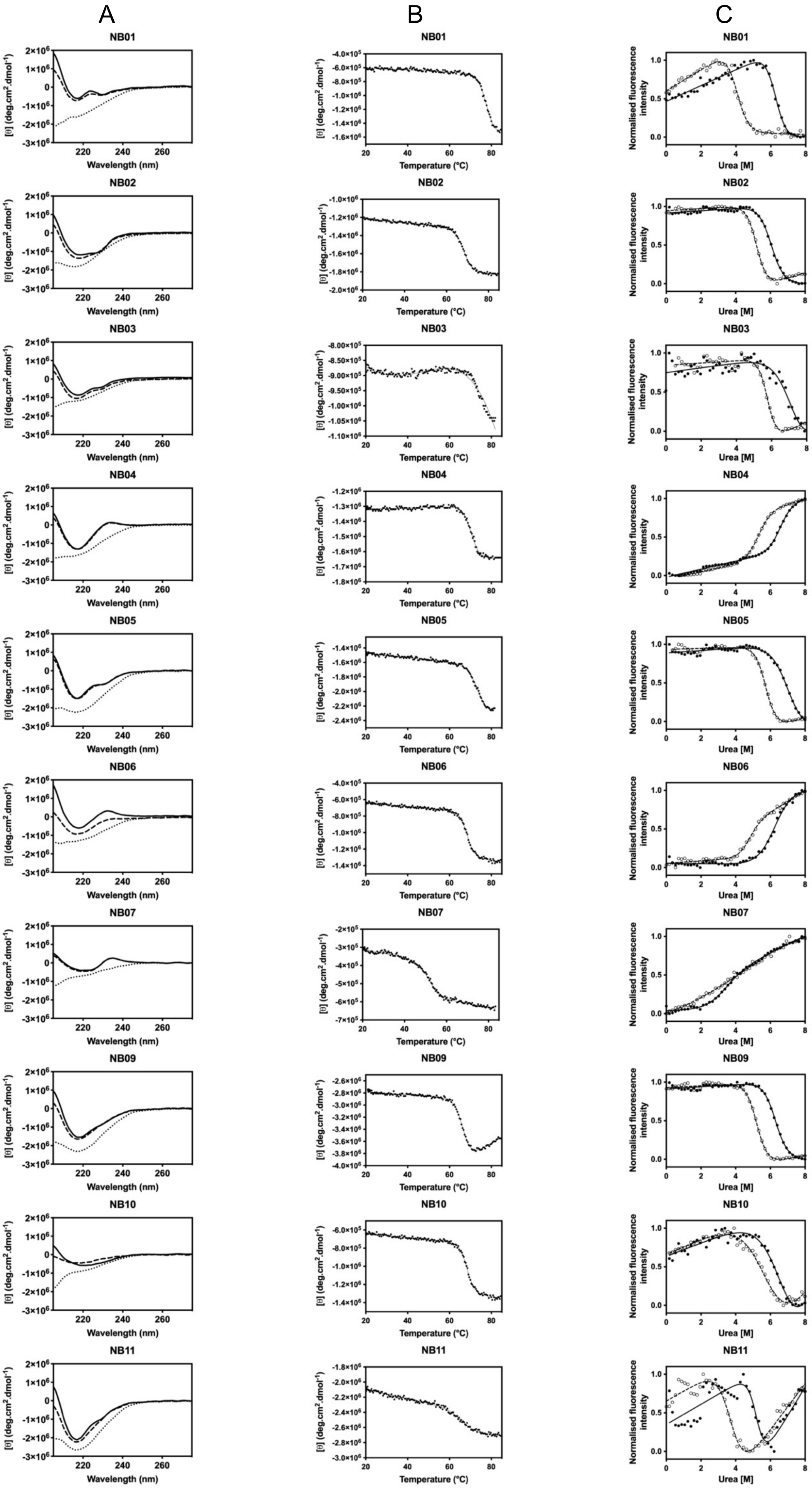
A
B
C
NB01
NB02
NB03
NB04
NB05
NB06
NB07
NB09
NB10
NB11
[θ] (deg.cm².dmol⁻¹)
Wavelength (nm)
Temperature (°C)
Normalised fluorescence intensity
Urea [M]

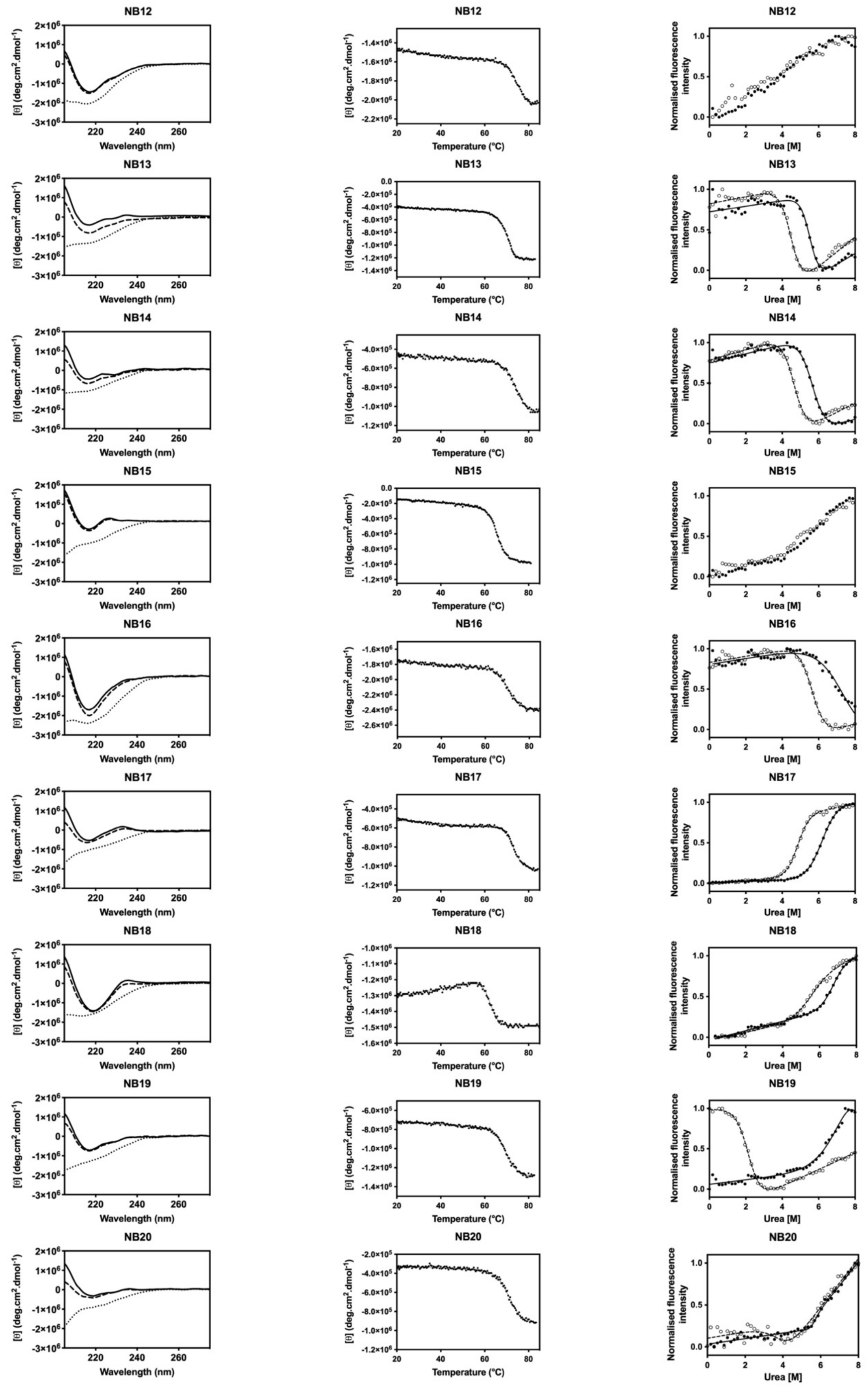


**Figure S2**. **Nanobody characterization.** Related to Figure 2. (A) Far-UV circular dichroism (far-UV CD) spectra, (B) thermal stability, (C) chemical denaturation monitored by tryptophan fluorescence. Of the 19 nanobodies tested, 15 showed a cooperative two-state unfolding. Nanobody NB08 did not express. Nanobodies NB06 and NB09 are included for comparison.

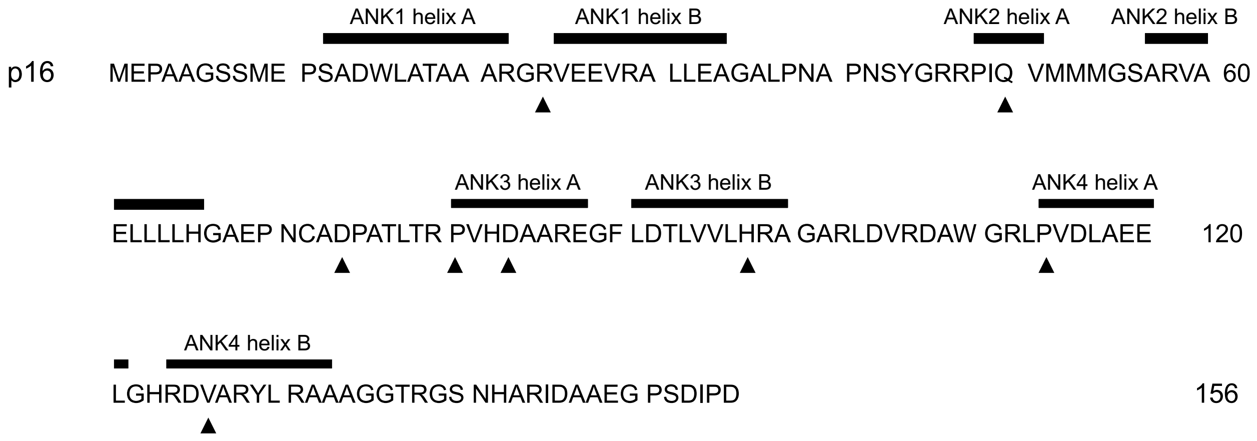


**Figure S3. Sequence of p16.** Related to Figure 4. The positions of the secondary structural ankyrin repeat elements are marked above the sequence. Black triangles identify the residues mutated within the set of p16 mutant proteins characterized in this study. The residues cover a range of different mutations previously identified in several different cancer types and are spread throughout p16. These mutations demonstrate a range of characteristics in comparison to p16 wildtype including increased aggregation propensity, compromised folding or temperature-dependent binding properties as previously shown (Parry and Peters, 1996; Zhang and Peng, 1996; Tang *et al.*, 1999; Hallett *et al.*, 2017) and further characterized here.

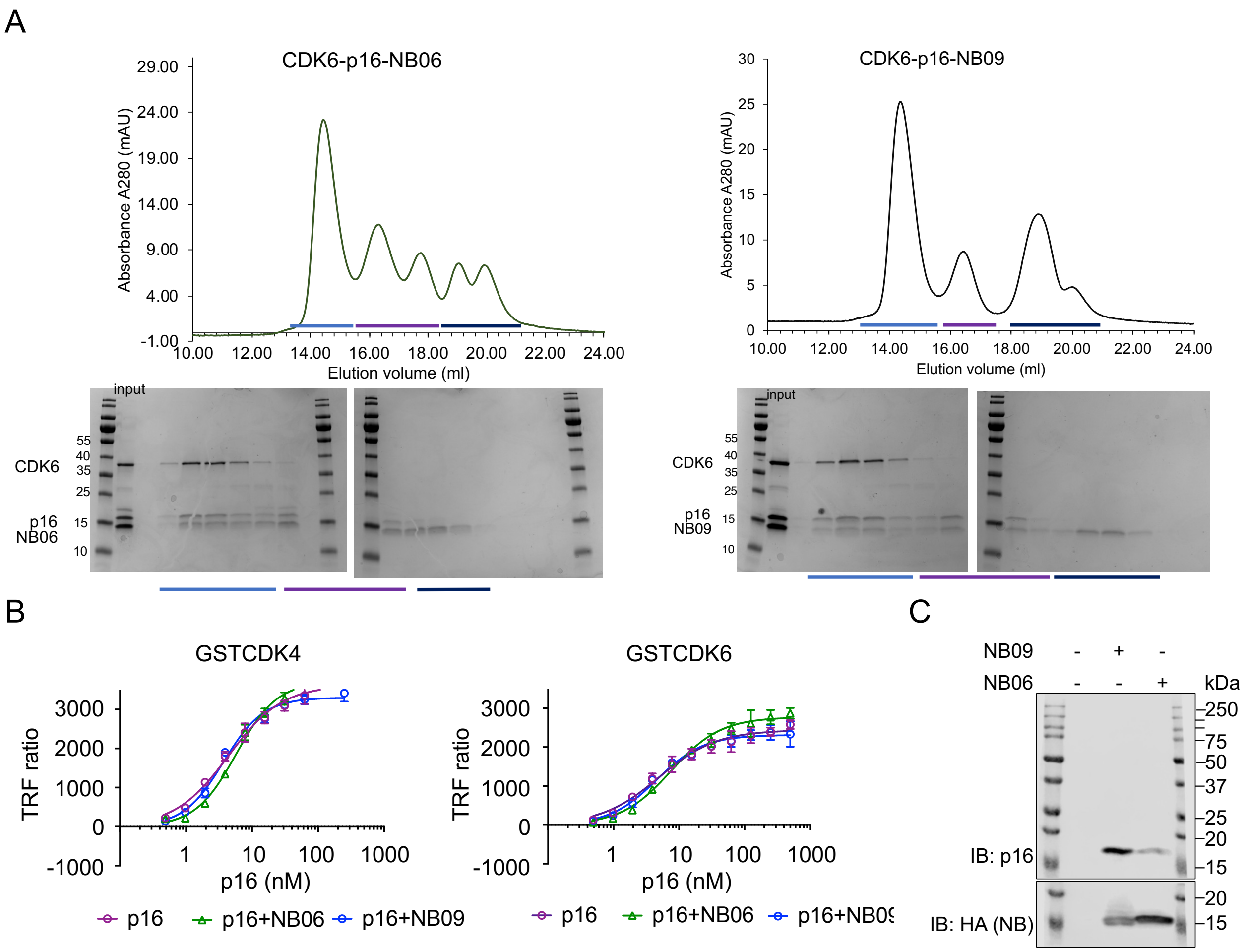


**Figure S4. Nanobodies NB06 and NB09 form a ternary complex with CDK6 and p16.** Related to Figure 5. (A) Nanobodies NB06 (green (LHS chromatogram) and NB09 (black (RHS chromatogram)) form a stable complex with CDK6 and p16. Nanobodies, p16 and CDK6 were purified separately. Chromatograms are to the same scale. Accompanying SDS-PAGE analysis of selected size-exclusion chromatography fractions is shown on the right. Samples were visualized by Instant Blue staining. The molecular weights of CDK6, p16, NB06 and NB09 are respectively 36 kDa, 16 kDa, and 14 kDa. Chromatogram is representative of two replicates carried out using independently prepared proteins. (B) CDK4 or CDK6 binding to p16 is not affected by presence of either NB06 or NB09 nanobodies as measured by homogenous time resolved fluorescence. The concentration of CDK4 and CDK6 used in these assays was 10 nM. (C) Nanobodies bind endogenous p16 in HEK293T cells. HEK293T cells were transfected with 5 µg of a plasmid containing either NB09 or NB06. Following a 24 hr incubation, cells were lysed, co-immunoprecipitated at 4°C for 4 hr with anti-HA beads and analyzed by western blot. Uncropped gel to accompany Figure 5D.

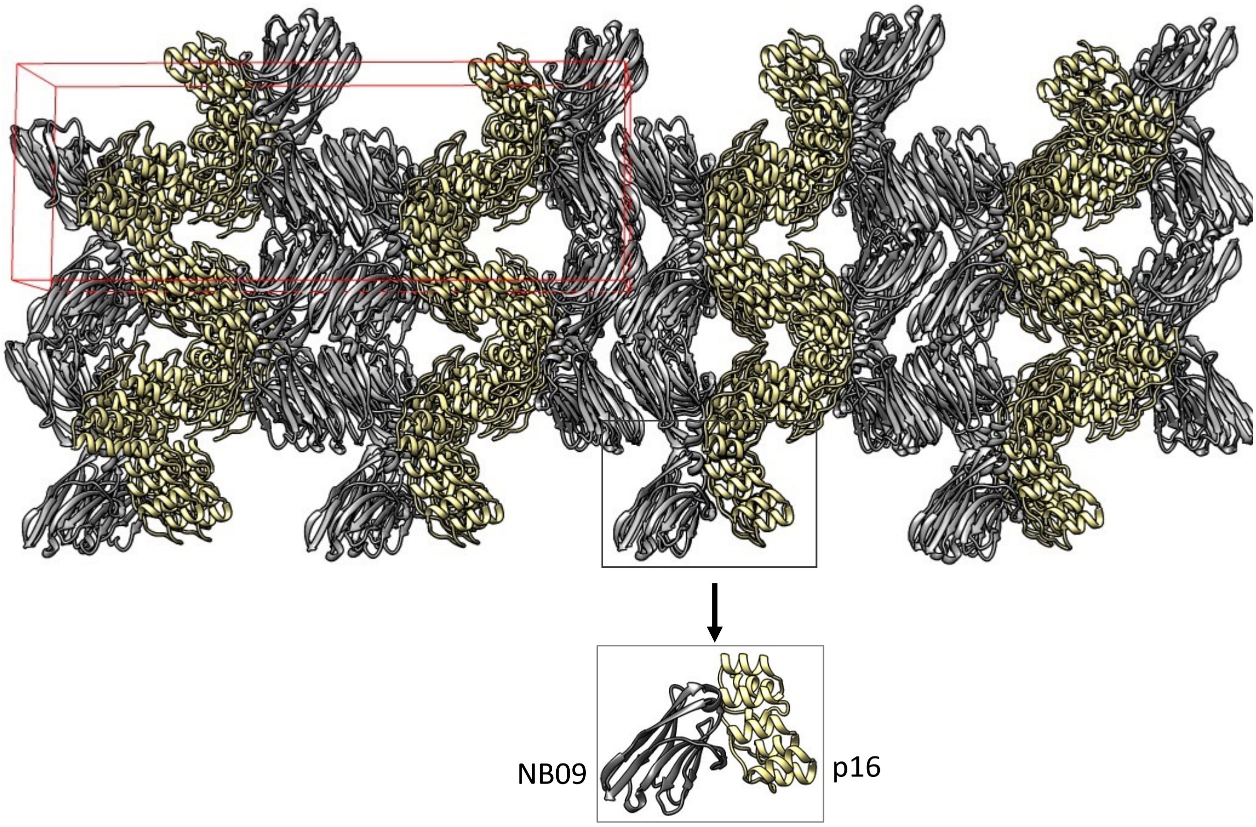


**Figure S5**. **p16-nanobody lattice.** Related to Figure 6. The location of one unit cell within the lattice is outlined in red. NB09 and p16 are rendered in ribbon representation in grey and straw respectively.

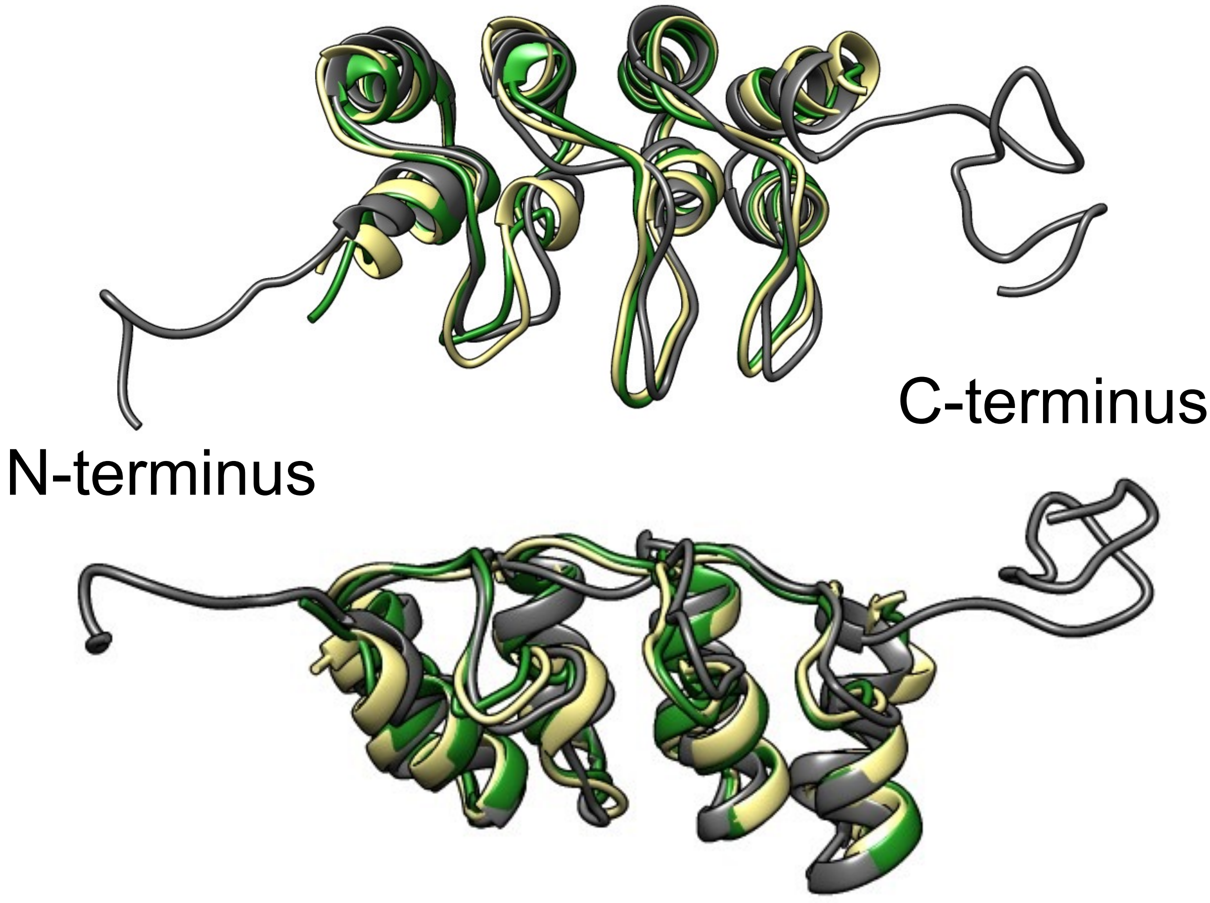


**Figure S6. Comparison of p16 structures.** Related to Figure 6. Superimposed structures of p16 from the p16-NB09 complex (straw), p16-CDK6 complex (green, PDB entry 1BI7) and p16 monomer structure determined by NMR (grey, PDB entry 2A5E). The upper and lower views are related by a rotation of 90$^{o}$ around a horizontal-axis.

**Table S1. Nanobody characterization.** Melting temperatures (Tm), and midpoints of unfolding (D50%) and m-values, were obtained by two-state fits of the thermal unfolding curves and chemical denaturation curves, respectively. Related to Figure 2.

| Nanobody | Tm (°C) | Without DTT D50% (M) | Without DTT *m*-value (kcal $mol^{-1}$ $M^{-1}$) | + 1 mM DTT D50% (M) | + 1 mM DTT *m*-value (kcal $mol^{-1}$ $M^{-1}$) |
|---|---|---|---|---|---|
| NB01 | 78.10 ± 0.2 | 6.26 ± 0.08 | 1.94 ± 0.22 | 4.03 ± 0.04 | 1.83 ± 0.17 |
| NB02 | 68.10 ± 0.0 | 6.04 ± 0.05 | 1.67 ± 0.12 | 5.22 ± 0.02 | 2.01 ± 0.09 |
| NB03 | 66.30 ± 1.2 | 7.32 ± 0.84 | 1.13 ± 0.31 | 5.78 ± 0.05 | 3.30 ± 0.32 |
| NB04 | 70.30 ± 0.1 | 6.56 ± 0.07 | 1.69 ± 0.13 | 5.32 ± 0.02 | 1.62 ± 0.07 |
| NB05 | 74.10 ± 0.2 | 7.10 ± 0.31 | 1.19 ± 0.13 | 5.74 ± 0.03 | 1.97 ± 0.12 |
| NB06 | 68.30 ± 0.0 | 6.24 ± 0.14 | 1.19 ± 0.13 | 4.72 ± 0.06 | 1.76 ± 0.25 |
| NB07[2] | 51.70 ± 0.1 | - | - | - | - |
| NB08[2] | - | - | - | - | - |
| NB09 | 66.60 ± 0.0 | 6.27 ± 0.05 | 1.71 ± 0.13 | 5.22 ± 0.02 | 1.90 ± 0.11 |
| NB10 | 68.30 ± 0.0 | 6.92 ± 0.42 | 0.90 ± 0.12 | 5.86 ± 0.20 | 0.81 ± 0.09 |
| NB11 | 65.60 ± 0.3 | 5.22 ± 0.10 | 2.26 ± 0.67 | 3.74 ± 0.06 | 1.49 ± 0.19 |
| NB12[2] | 75.20 ± 0.2 | - | - | - | - |
| NB13 | 70.60 ± 0.1 | 5.53 ± 0.07 | 2.32 ± 0.45 | 4.50 ± 0.04 | 1.93 ± 0.19 |
| NB14 | 75.00 ± 0.2 | 5.66 ± 0.07 | 1.63 ± 0.21 | 4.66 ± 0.03 | 1.79 ± 0.12 |
| NB15[2] | 65.50 ± 0.1 | - | - | - | - |
| NB16 | 70.20 ± 0.1 | 7.51 ± 0.36 | 0.82 ± 0.12 | 5.64 ± 0.05 | 1.67 ± 0.17 |
| NB17[1] | 72.70 ± 0.1 | 6.19 ± 0.02 | 1.39 ± 0.03 | 4.84 ± 0.02 | 1.86 ± 0.09 |
| NB18 | 62.50 ± 0.1 | 6.78 ± 0.12 | 1.64 ± 0.18 | 5.45 ± 0.16 | 1.15 ± 0.19 |
| NB19[1] | 69.70 ± 0.1 | 8.08 ± 0.77 | 0.93 ± 0.11 | 2.16 ± 0.03 | 1.76 ± 0.09 |
| NB20 | 71.80 ± 0.2 | 5.60 ± 0.14 | 3.99 ± 2.90 | 4.02 ± 0.25 | 1.40 ± 0.59 |

[1] NB17 and NB19 contain two disulphide bonds and respond differently: The loss of the disulphide bonds in NB19 had a very large effect on its stability. This nanobody contains the second longest CDR3 loop of all the nanobodies tested, which is stabilized by a second disulphide bond. In contrast, the stability of NB17 was less affected by their loss.

[2] Four of the nanobodies showed no discernible unfolding transition with or without the disulphide bond, presumably a result of having very high stabilities.

**Table S2.** Nanobodies stabilize p16. Related to Figure 3.

| p16/nanobody | $D_{50\%}$ (M)[1] | $m$ (kcal mol$^{-1}$ M$^{-1}$ ) | $\Delta G_{unf}$ (kcal mol$^{-1}$)[2] |
|---|---|---|---|
| p16 | 1.94 ± 0.01 | 1.66 ± 0.02 | 3.22 ± 0.04 |
| p16 + NB03 | 1.94 ± 0.01 | 1.61 ± 0.03 | 3.22 ± 0.04 |
| p16 + NB05 | 2.04 ± 0.02 | 1.72 ± 0.07 | 3.39 ± 0.05 |
| p16 + NB06 | 2.80 ± 0.01 | 1.58 ± 0.03 | 4.65 ± 0.06 |
| p16 + NB09 | 3.11 ± 0.04 | 1.35 ± 0.05 | 5.16 ± 0.09 |
| p16 + NB16 | 1.92 ± 0.03 | 1.67 ± 0.07 | 3.19 ± 0.06 |
| p16 + NB17 | 2.01 ± 0.01 | 1.45 ± 0.03 | 3.34 ± 0.04 |

[1] Denaturation experiments were performed at 25 °C in 50 mM Tris-HCl pH 7.5, 100 mM NaCl. The concentration of each protein was 2 µM. Data were fitted to a two-state model to obtain the midpoint of unfolding, $D_{50\%}$, and the *m*-value. N=3 ± SD.

[2] The free energy of unfolding, $\Delta G_{unf}$, was calculated as the product of the midpoint of unfolding, $D_{50\%}$, and the *m*-value for wild-type p16 (in the absence of any nanobody).

**Table S3. X-ray data collection and refinement statistics.** Related to Figure 6.

| | |
|---|---|
| Complex | p16-NB09 |
| BAG number | mx13587-74 |
| Beamline | I03 |
| PDB code | 7OZT |
| **Data collection** | |
| Space Group | C 2 2 $2_1$ |
| Unit Cell dimensions (a,b,c) (Å) | 42.12, 182.93, 65.23 |
| Unit Cell dimensions (α,β,γ) (°) | 90.00, 90.00, 90.00 |
| Resolution limits (Highest resolution shell) (Å) | 91.41 – 1.74 (1.77 – 1.74) |
| Observations | 180628 (8689) |
| Unique Reflections | 26446 (1412) |
| Completeness (%) | 100 (100) |
| $R_{merge}$[1] | 0.108 (0.878) |
| **Refinement** | |
| Number of atoms | |
| Protein | 1857 |
| Water | 94 |
| Resolution (Å) | 91.47-1.74 (1.77-1.74) |
| Reflections | 26427 (930) |
| $R_{cryst}$ /$R_{free}$[2] | 0.188 / 0.217 |
| R.m.s. deviations | |
| Bond lengths (Å) | 0.0122 |
| Bond angles (°) | 1.84 |
| B-factors (Å$^2$) | |
| Protein | 29.8 |
| Water | 32.8 |

[1] $R_{merge} = \Sigma_h\Sigma_i|I_{h,i} - \langle I_h\rangle| / \Sigma_h\Sigma_i I_{h,i}$.

[2] $R_{cryst} = \Sigma h||F_{o,h}| - |F_{c,h}|| / \Sigma|F_{o,h}|$, where $F_o$ and $F_c$ are the observed and calculated structure factors. $R_{free}$ was determined from 5% data.

**Table S4. Oligonucleotide sequences.** Related to key resources table. Sequences in lower case are primer overhangs used for InFusion cloning, sequences in upper case anneal to the p16 or NB06/NB09 coding regions as indicated and are used for PCR amplification.

| **Oligonucleotides** | | |
|---|---|---|
| MP57 forward | 5'-TTATGCTTCCGGCTCGTATG-3' | Nanobody pMESy4 forward sequencing primer |
| GIII | 5'-CCACAGACAGCCCTCATAG-3' | Nanobody pMESy4 reverse sequencing primer |
| T7 forward | 5'-TAATACGACTCACTATAGGG-3' | T7 forward sequencing primer |
| TriexDown reverse | 5'-TCGATCTCAGTGGTATTTGTG-3' | pOPIN reverse sequencing primer |
| p16 Forward | 5'- aagttctgtttcagggCCCGATGGAGCC -3' | Amplifies p16 |
| p16 Reverse | 5'- atggtctagaaagctttaATCGGGGATGTCTGAGGGAC -3' | Amplified p16 |
| $p16^{NTA}$ Tag Forward | 5'-aagttctgtttcagggcccgGGCCTGAACGATATCTTCGAAG - 3' | Amplifies N-terminal AviTag and adds handles for InFusion cloning to vector |
| $p16^{NTA}$ Tag Reverse | 5'-cggctccatTTCATGCCATTCAATCTTCTGC -3' | Amplifies N-terminal AviTag and adds handles for InFusion cloning to p16 |
| $p16^{NTA}$ Protein Forward | 5'-ggcatgaaATGGAGCCGGCGGCG-3' | Amplifies p16 and adds handles for InFusion cloning to N-terminal AviTag |
| $p16^{NTA}$ Protein Reverse | 5'- atggtctagaaagctttaATCGGGGATGTCTGAGGGAC-3' | Amplifies p16 and adds handles for InFusion cloning to vector. |
| $p16^{CTA}$ Tag Forward | 5'- tccccgatGGCCTGAACGATATCTTCGAA-3' | Amplifies C-terminal AviTag and adds handles for InFusion cloning to p16. |
| $p16^{CTA}$ Tag Reverse | 5'-atggtctagaaagctttattaTTCATGCCATTCAATCTTCTGC-3' | Amplifies C-terminal AviTag and adds handles for InFusion cloning to vector. |
| $p16^{CTA}$ Protein Forward | 5'- aagttctgtttcagggCCCGATGGAGCC -3' | Amplifies p16 and adds handles for InFusion cloning to vector. |
| $p16^{CTA}$ Protein Reverse | 5'-ttcaggccATCGGGGATGTCTGAGGG -3' | Amplifies p16 and adds handles for InFusion cloning to C-terminal AviTag |
| NB06 Fwd | 5'- tcaaaggagatatac CCATGGCCCAGGTTCAGC | Amplifies NB06 and adds handle for InFusion cloning into pOPINF-HA |

| | | |
|---|---|---|
| NB06-Rev | 5’-cacgtcgtaggggtagaattcGCTGCTAACGGTAACCTG | Amplifies NB06 and adds handle for InFusion cloning into pOPINF-HA |
| NB09-Fwd | 5’- tcaaaggagatatacCCATGGCCCAGGTTCAGC | Amplifies NB09 and adds handle for InFusion cloning into pOPINF-HA |
| NB09-Rev | 5’- cacgtcgtaggggtagaattcACTGGAGACGGTGACCTG | Amplifies NB09 and adds handle for InFusion cloning into pOPINF-HA |
